# Flexible and scalable control of T cell memory by a reversible epigenetic switch

**DOI:** 10.1101/2022.12.31.521782

**Authors:** Kathleen Abadie, Elisa C. Clark, Obinna Ukogu, Wei Yang, Riza M. Daza, Kenneth K.H. Ng, Jumana Fathima, Allan L. Wang, Avinash Bhandoola, Armita Nourmohammad, Jay Shendure, Junyue Cao, Hao Yuan Kueh

**Author notes:** These authors contributed equally to this manuscript (Kathleen Abadie, Elisa C. Clark). These authors contributed equally to this manuscript (Jay Shendure, Junyue Cao, Hao Yuan Kueh).

## Abstract

The immune system encodes information about the severity of a pathogenic threat in the quantity and type of memory cell populations formed in response. This encoding emerges from the decisions of lymphocytes to maintain or lose self-renewal and memory potential during a challenge. By tracking responding CD8 T cells at the single-cell and clonal lineage level using time-resolved transcriptomics and quantitative live imaging, we identify a remarkably flexible decision-making strategy, whereby T cells initially choose whether to maintain or lose memory potential early after antigen recognition, but following pathogen clearance may regain memory potential if initially lost. Mechanistically, this flexibility is implemented by a *cis*-epigenetic switch that silences the memory regulator TCF1 in a stochastic and reversible manner in response to stimulatory inputs. Mathematical modeling shows how this strategy allows memory T cell numbers to scale robustly with pathogen virulence and immune response magnitudes. We propose that flexibility and stochasticity in cellular decision making ensures optimal immune responses against diverse threats.

## Main

The immune system keeps a memory of prior infections with information about the inducing threat. This memory is encoded by the numbers and types of memory lymphocytes generated upon challenge. The quantity of memory T cells, in particular, scales with the magnitude of a prior infection, such that the memory population is a fixed fraction of the T cell number at the infection peak, across a range of pathogenic challenges^1–3^. This scaling in memory production is robust across T cell clones with different epitope specificities and allows the body to generate memory proportional to the severity of the pathogenic challenge. The regulatory mechanisms that enable this critical feature of adaptive immunity are not well understood.

The size and characteristics of the memory compartment are determined by the fate decision-making strategies of T cells responding to an acute infection^4^. As naive CD8 T cells respond to antigens, they must decide whether and when to maintain long-term viability and self-renewal potential, and thereby persist to form memory cells as the infection is cleared. One class of models posits that cells make this decision early after antigen encounter, and in a mutually exclusive manner with effector differentiation (Fig. 1A)^5–7^. Under this model, memory cells form directly from naive cells without first passing through an effector phase, but through an early lineage bifurcation that concurrently gives rise to short-lived effector cells. A second class of models posits that cells decide later, only after they have undergone effector differentiation (Fig. 1A)^8–10^. In this strategy, cytotoxic effectors that maintain memory potential populate the memory compartment upon infection clearance. However, in contrast with both models, it is also possible that this process is inherently flexible^11^, such that T cells have multiple opportunities to commit to the memory state. From a social and cognitive sciences perspective^12,13^, flexibility in decision making allows individuals to adapt and better respond to uncertain and dynamic environments; in the immune system, such flexibility may allow T cells to optimize memory formation for threats whose properties might only manifest as they unfold in time. It is unclear whether there exists such flexibility in T cell memory formation and, if so, what its underlying mechanisms and functional roles are.

**Figure 1:**
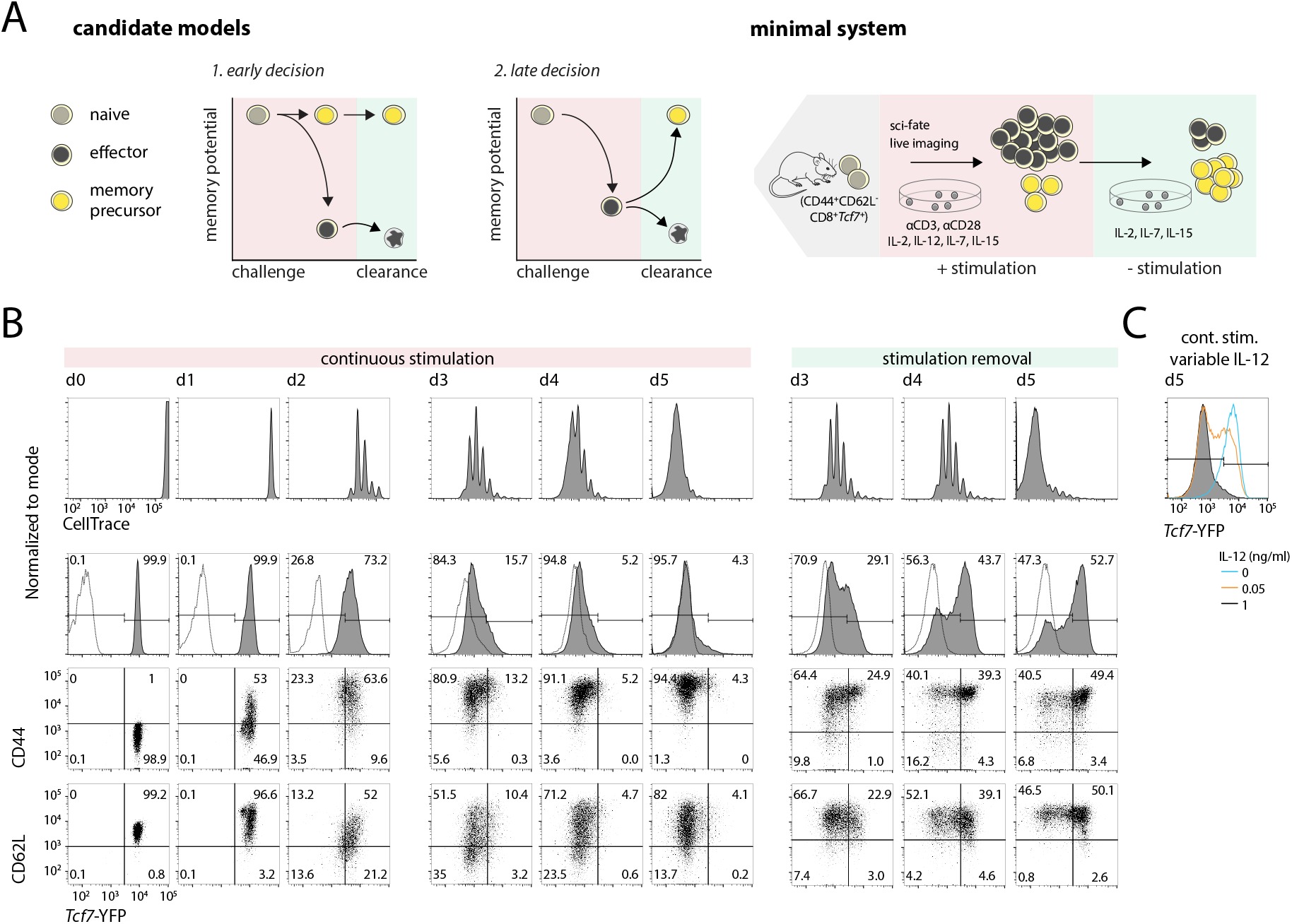
A minimal *ex vivo* system to track CD8 T cell effector and memory decision making dynamics. (**A**) Candidate decision-making strategies for CD8 T cell memory generation (left); a minimal *ex vivo* system for tracking memory decision-making dynamics at the single-cell level. (**B**-**C**) Naive CD8 T cells were isolated from *Tcf7*-YFP reporter mice, then cultured using this minimal *ex vivo* system. Flow cytometry plots show analysis of cultured CD8 T cells during initial stimulation for 2 days (left) and continued stimulation to day 5 (middle), or after stimulation withdrawal (removal of αCD3/αCD28 after day 2 and IL-12 after day 3) (right). (**C**) *Tcf7*-YFP silencing is tunable by IL-12 levels. Data are from a single experiment representative of at least 3 independent experiments.

In this study, we sought to elucidate the memory decision-making dynamics of CD8 T cells by following the regulation of TCF1 (encoded by *Tcf7*), a transcription factor essential for memory cell generation^14^. *Tcf7* is expressed in naive and memory cells, where it is crucial for maintaining self-renewal, and is silenced during effector differentiation, resulting in loss of memory potential and entry into a short-lived state^5,15^. To follow *Tcf7* regulation and memory decision-making in a controlled environment where cells can be continuously observed and signaling inputs carefully manipulated, we developed an *ex vivo* system to mimic stimulation of T cells by acute challenge. Using this system and complementary testing *in vivo*, we uncover a flexible decision-making strategy: T cells can gain or lose memory potential at multiple junctures after antigen encounter, and do so in a stochastic and reversible manner. Mathematical modeling reveals that this flexible decision-making strategy allows for the number of memory cells to scale linearly with total numbers of expanded T cells at the peak of infection, thereby encoding information about the severity of the prior threat.

### A minimal *ex vivo* system for effector and memory differentiation

In our system, naive (CD44^-^CD62L^+^) CD8 T cells with a knock-in YFP reporter for *Tcf7*^16^ are activated with plate-immobilized anti-CD3 and anti-CD28 antibodies and IL-2, together with additional cytokines present during acute infection (IL-12, IL-7, and IL-15^17–19^). These conditions minimize variability in the exposure of individual cells to stimulatory signals, enabling cell-intrinsic lineage control mechanisms to be studied apart from environmental heterogeneity.

In this system, all cells begin dividing rapidly after 24 hours and upregulate the transmembrane glycoprotein CD44, indicating uniform activation (Fig. 1B). Activated cells downregulate *Tcf7* and the lymph node-homing adhesion molecule CD62L, consistent with effector differentiation. The inflammatory cytokines IL-12 and IFN-β1 enhance *Tcf7*-YFP silencing (Fig. 1C, Fig. S1C-D), consistent with their roles in driving effector differentiation^20,21^. When TCR stimulation (anti-CD3/CD28) and inflammation (IL-12) are removed to mimic pathogen clearance, the cells demonstrate a population-level increase in CD62L and *Tcf7*-YFP while continuing to divide, as previously observed^4^. *Tcf7* and CD62L levels are heterogeneous both during stimulation and after removal, suggestive of an early memory and effector differentiation decision. YFP levels closely matched TCF1 protein levels throughout activation, validating use of the reporter in this system (Fig. S1A-B).

### Naive cells bifurcate early into effectors and memory precursors

To determine whether the heterogeneity in *Tcf7* and CD62L downregulation reflects early memory and effector programming (Fig. 1), we analyzed *ex vivo* activated cells using the temporally-resolved single-cell transcriptome sequencing method, *sci-fate*^22^. Here, metabolic labeling of newly-synthesized transcripts reveals a cell’s current activity state apart from its history^22,23^ (Fig. 2A). We subjected cells at days 1, 2, and 4 to 4-thiouridine (4sU) pulse-labeling for 2 hrs, followed by sequencing and analysis as previously described^22^. We obtained old and new transcriptomes for ~17,000 single cells, with a median of 17,574 total and 2,529 new transcripts detected per cell (Fig. S2A). To disentangle effector and memory gene programs from other activation-induced programs, we performed an integrative analysis of our temporally-resolved transcriptome data and existing transcription factor (TF) binding data^24^ to identify TF modules, consisting of co-regulated groups of TFs and their cognate target genes (see Methods). This analysis revealed two main TF modules, a cell cycle module and a T cell differentiation module, the latter further separable into submodules that included known regulators of effector and memory differentiation (Fig. 2B; Fig. S2C).

**Figure 2:**
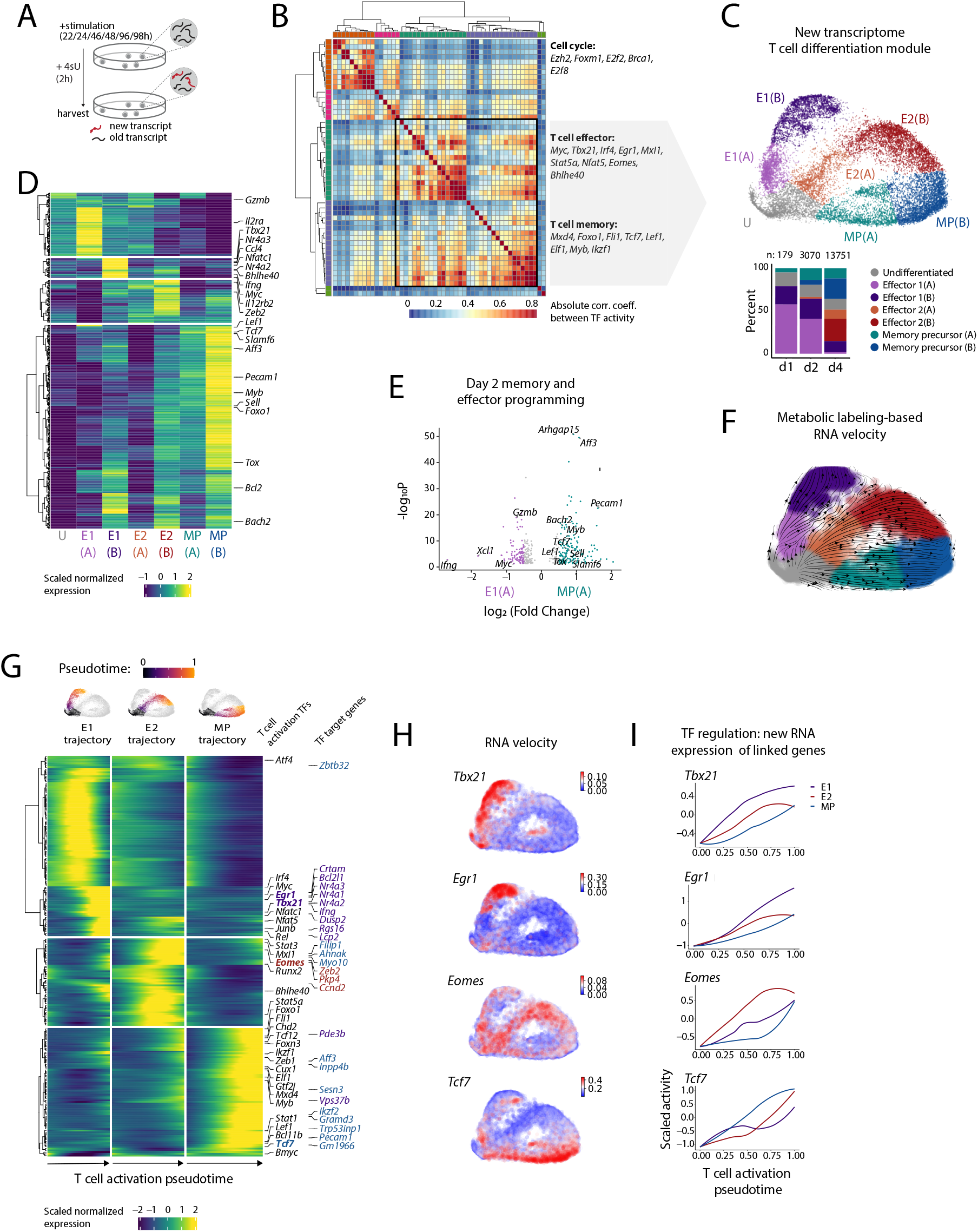
Naive cells diverge into effector and memory states early after activation. (**A**) Naive CD8 T cells were activated as in Fig. 1A, with 0.05 ng/ml IL-12. After 1, 2, and 4 days, cells were treated with 4sU for 2 hours to label new transcripts, then harvested for time-resolved transcriptomics using *sci-fate*. (**B**) Heatmap showing the absolute Pearson’s correlation coefficient between the activities of pairs of TFs, generated using *sci-fate*. Key TFs in each module are labeled at right. T cell differentiation module used for subsequent analysis is boxed. (**C**) UMAP visualization of cells based on the activity of T cell differentiation-related TF module, using newly synthesized mRNA, colored by cluster ID (top). Percentage of cells in each T cell activation state cluster after indicated days (bottom). (**D**) Aggregated expression (scaled, log_10_ normalized) of top 400 differentially expressed (DE) genes between clusters (q < 3 x 10^-45^ for all genes except for *Ifng*, q = 7.3 x 10^-29^). (**E**) DE genes between E1(A) and MP(A) at day 2 only; log2FC > 0.5 and adj. p < 0.05. (**F**) UMAP visualization as in (C), characterized by labeling-based RNA velocity analysis. Streamlines indicate the integration paths that connect local projections from the observed state to the extrapolated future state(*26*). (**G**) Pseudotemporal ordering of top 200 DE genes and additional genes of interest (q < 1.4 x 10^-17^) between trajectories. Gene labels correspond to all DE TFs in the T cell differentiation TF module (left text) and DE target genes linked to *Tbx21, Egr1, Eomes*, and *Tcf7* (right text). (**H**) RNA velocity and (**I**) Loess smoothed TF activity over pseudotime for four of the most DE genes between trajectories. TF activity is calculated as the normalized aggregation of levels of newly synthesized mRNA for all TF target genes, scaled across all cells. Cells in the undifferentiated (U) cluster are set to pseudotime = 0 for each trajectory.

By visualizing cell states using genes in the T cell differentiation module for Uniform Manifold Approximation and Projection (UMAP) dimensionality reduction, we resolved distinct effector and memory states with coherence between timepoints (Fig. 2C; Fig. S2B, D). Unsupervised clustering and differential gene expression analysis revealed distinct early and late (A vs. B) effector (E1 and E2) and memory precursor (MP) states. E1 and E2 cells exhibited higher expression of the effector-associated genes *Gzmb, Ifng, Tbx21, Zeb2*, and *IL12rb2*, while MP cells had higher expression of the stem- and memory-associated factors *Bach2, Lef1, Tcf7, Sell*, and *Slamf6*, and lower expression of effector-associated genes (Fig. 2D-E; Fig. S2E-F; Supplementary Table 1-2)^25^. These differential gene expression patterns were present at day 2 and amplified at day 4.

Consistent with an early fate bifurcation, RNA velocity vectors calculated using reads from newly synthesized transcripts originate from the undifferentiated state (U), and flow along separate effector and memory branches^26^ (Fig. 2F). To gain insight into the dynamics of genes differentially regulated between divergent trajectories, we visualized their expression over pseudotime along each trajectory (Fig. 2G; Supplementary Table 3). This analysis, together with RNA velocity and TF activity analysis (Fig. 2H-I), identified effector and memory regulators with greatest differential regulation along their respective trajectories. *Tbx21, Egr1*, and *Irf4*,among other effector regulatory genes, were specifically active along the E1 trajectory, while a distinct set of effector regulators, including *Eomes, Bhlhe40, Stat5a* and *Stat3*, characterize the E2 trajectory. This effector heterogeneity and its potential influence on downstream differentiation will be interesting to investigate in future studies but is not further pursued here. Finally, regulators of T cell stemness and survival, including *Tcf7, Myb, Mxd4* and *Fli1*, were active in the MP trajectory. *Tcf7* was the most significantly differentially expressed gene between trajectories, upregulated early along the MP trajectory and absent in both E1 and E2 trajectories. Its expression furthermore coincided with that of target genes identified through TF linkage that promote self-renewal, such as *Ikzf2, Sesn3, Aff3*, and *Pecam1* (CD31). Thus, *Tcf7* is a critical driver of this early divergent memory trajectory in our system.

### The early effector and memory decision occurs heterogeneously within clones

The divergence of cells into effector and memory lineages, occurring even under the strong, uniform stimulatory conditions of our *ex vivo* system, is suggestive of a cell-intrinsic regulatory mechanism involving *Tcf7* that generates heterogeneity in fate outcomes. To elucidate the degree to which this decision is heterogeneous within cell lineages amid constant environmental signals, we acquired multi-day time-lapse movies of clonal CD8 T cell lineages during activation with continuous measurement of *Tcf7* reporter levels (Fig. 3). As T cells are difficult to track with live imaging due to their high mobility, tendency to adhere to one another, and rapid proliferation, we optimized adhesion conditions and computational analyses that allow continuous tracking of a fate regulating TF across clonal CD8 T cell lineages (Fig. 3; Fig. S3; see Methods)^27^. Using this method, we tracked a total of 120 clonal lineages over 4 days and an average of 4.4 cell generations.

**Figure 3:**
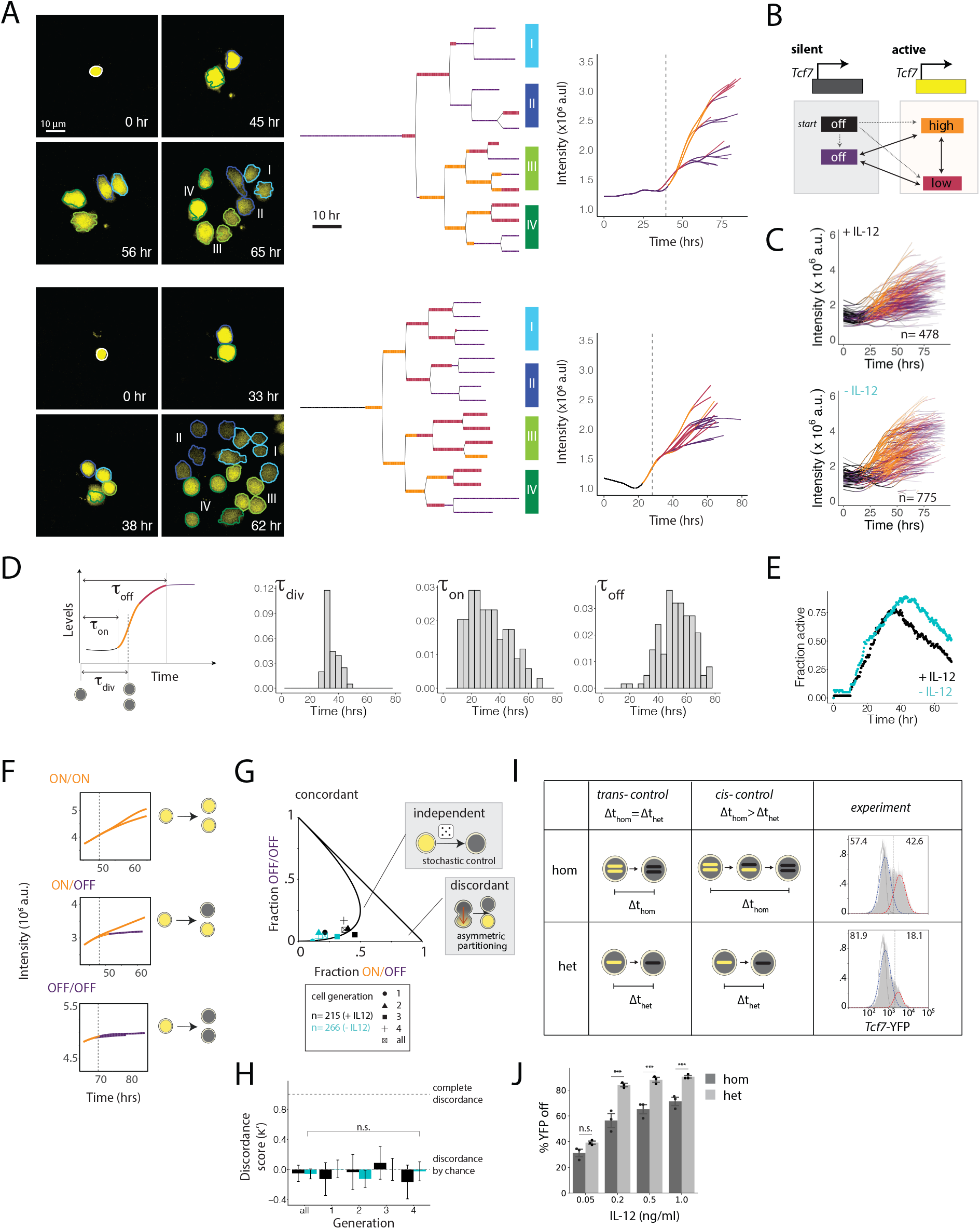
Heterogeneous *Tcf7* silencing within clones is controlled by a stochastic epigenetic switch. (**A**) Representative lineages demonstrating clonal heterogeneity in *Tcf7*-YFP silencing: image snap shots (left), lineage trees (middle), and reporter intensity (area x median YFP levels) over time for each track (right), with the first cell division marked by a vertical dashed line. Cell borders in snapshots are colored and labeled to match their corresponding leaves in the lineage trees. Lineage trees and tracks are colored by HMM-derived promoter state, outlined in (**B**). Cells are cultured with 1 ng/ml IL-12 unless otherwise indicated. (**C**) Reporter intensity for all overlaid tracks, colored by promoter state. (**D**) For each track, from left to right: time of first division, time of first transition to a stable active state, time of first transition to a stable silent state (stable state ≥ 10 hrs). (**E**) For all lineages combined, fraction of cells in an active promoter state over time, +/- 1 ng/ml IL-12. (**F-H**) Each division of a parent cell with the *Tcf7* promoter ON was categorized as giving rise to zero, one, or two daughters that transition to an OFF state. (**F**) Examples of each division category. (**G**) The OFF/OFF fraction by ON/OFF fraction is plotted separately for each generation, +/- IL-12, to distinguish concordant, independent, and asymmetric silencing mechanisms. (**H**) Modified Cohen’s kappa test for division events in (G). (**I**) Comparison of YFP/YFP and YFP/+ reporters to distinguish *cis* and *trans* regulation of *Tcf7* silencing (left). YFP distributions for YFP/YFP and YFP/+ reporters cultured for 5 days with 0.2 ng/ml IL-12 (right). YFP off fractions are calculated from gaussian fits to distributions. (**J**) YFP off percentages as in (I), over a range of IL-12 concentrations. Mean ± s.d. Statistical significance was calculated with an unpaired two-tailed t test; n.s. p=0.05, ***p<0.005. Individual data points are from a single experiment representative of 2 independent experiments (I-J).

Naive cells in these time-lapse movies start small, adhere to the antibody-bound plate, acquire CD69 expression, increase dramatically in size, and divide rapidly after 1-2 days (Fig. S3G; Supplemental movie 1). Strikingly, individual activating T cell clones often gave rise to *Tcf7* high and low subpopulations (Fig. 3A; Fig. S3J; Supplementary Movie 1), indicating that the effector and memory decision is made heterogeneously within clones. Of note, *Tcf7* low and high cells showed similar degrees of attachment to the surface, indicating that these intra-clonal differences are not due to differences in TCR stimulation, but more likely due to cell-intrinsic mechanisms generating heterogeneity in *Tcf7* silencing.

Differences in *Tcf7*-YFP levels after multiple cell divisions likely stem from earlier *Tcf7* silencing events propagated through dilution of the stable fluorescent protein by cell division. To pinpoint the timing of early regulatory events that give rise to these differences in *Tcf7-* YFP levels, we calculated the *Tcf7* promoter activity over time in single cells, defined as the rate at which total *Tcf7-*YFP levels increase over time, using a Hidden Markov Model (HMM) to assign *Tcf7* promoter activity states to each cell at each timepoint and identify switching points between those states (Fig. 3A-C; Fig. S3A-F; see Methods).

This analysis revealed that cells silence *Tcf7* expression at variable times after the onset of stimulation, and can do so as early as the first cell division as well as at later generations. Cells activated the *Tcf7* promoter prior to the first cell division, reflecting exit from quiescence, and then proceeded to switch the *Tcf7* promoter to a silent state. The timing at which the *Tcf7* promoter transitioned to the silent state varied between cell tracks both within and between cell lineages, consistent with observed heterogeneity in *Tcf7*-YFP levels within clones (Fig. 3A-D). Removing IL-12 increased the fraction of cells in an active promoter state (Fig. 3C,E). Silent *Tcf7* promoter states persisted across multiple cell divisions (Fig. 3A; Fig. S3I-J) and thus represent heritable regulatory changes as opposed to more transient dynamics such as transcriptional bursting. These results provide evidence that a cell-intrinsic *Tcf7* silencing event, occurring heterogeneously within clones, underlies the early divergence in effector and memory states.

### A stochastic epigenetic switch controlling *Tcf7* silencing underlies the early effector and memory decision

Heterogeneity in *Tcf7* silencing could derive from asymmetric cell division^6,28^, whereby cell fate determinants partition unequally, giving rise to discordant behavior between two sister cells. Alternatively, this heterogeneity could result from stochastic control^29–31^, whereby two sisters would make *Tcf7* silencing decisions independently. While two sister cells could still make different decisions, they would silence *Tcf7* discordantly no more frequently than expected by chance. To test these predictions, we analyzed the fractions of daughter cell pairs that silence *Tcf7* either discordantly (ON/OFF) or concordantly (OFF/OFF), doing so for cell pairs across all cell generations, with or without IL-12 (Fig. 3F). By plotting concordant (OFF/OFF) versus discordant (ON/OFF) sister pair fractions, we found that all data points adhered to a theoretical curve representing the expected relationship between sister pair fractions for independent regulation (Fig. 3G). Consistently, by statistical analysis using a modified Cohen’s kappa coefficient (κ’), we found that daughter cells were no more likely to make discordant decisions than expected by chance (Fig. 3H; Supplementary Table 4). These findings support the view that *Tcf7* silences in a stochastic manner to drive divergent decision making within clones.

Epigenetic switching mechanisms, involving changes in chromatin modifications or conformation at gene loci, can introduce stochastic rate-limiting steps to gene activation or silencing^32–34^. As *Tcf7* silencing involves repressive DNA or histone methylation^14,20,35,36^, it could be gated by such a mechanism. Epigenetic mechanisms act in *cis* at individual gene loci and therefore would silence each *Tcf7* locus independently. To test for this mechanism, we compared *Tcf7-*YFP silencing kinetics in cells from mice homozygous and heterozygous for the reporter, with the prediction that homozygous reporter cells would yield a smaller population of YFP-low cells, since both loci need to silence for loss of reporter expression (Fig. 3I-J, Fig. S3K). Indeed, the *Tcf7*-YFP silent population was significantly smaller in homozygous reporter cells and increased with IL-12, consistent with a *cis*-epigenetic silencing mechanism modulated by inflammation. Together, these results provide evidence that a stochastic *cis*-epigenetic switch, tunable by external stimuli, controls the early decision of naive cells to silence *Tcf7* expression and memory potential.

### Reversibility of *Tcf7* silencing enables a late memory decision

*Tcf7* silencing has been proposed to be an irreversible event that marks a ‘point of no return’ for effector differentiation and loss of memory potential^5,37^. Conversely, various studies have demonstrated that cells that acquire cytotoxic effector function are able to populate memory compartments after an infection is resolved^8,9,38^, suggesting that *Tcf7*-silenced effectors may still be able to reactivate *Tcf7* and reacquire memory potential. Our data thus far provide evidence for an early T cell decision to abandon or maintain memory potential, driven by stochasticity in antigen-driven *Tcf7* silencing, but do not exclude the possibility that effector cells can reverse their decisions and regain memory potential later after withdrawal of stimulation.

To test this possibility, we sorted *Tcf7*-YFP low and *Tcf7*-YFP high cells after initial culture and subjected them to reculture with variable stimulation conditions *ex vivo* (Fig. 4A). As expected, sorted *Tcf7*-YFP high cells maintained *Tcf7-*YFP expression without stimulation but underwent heterogeneous silencing under continuing stimulation (Fig. 4B-C, Fig. S4A-B). Furthermore, *Tcf7*-YFP low cells maintained a silent state upon continued stimulation, as observed. Strikingly, however, upon stimulation withdrawal, *Tcf7* reactivated, with the fraction of *Tcf7* expressing cells increasing over 6 days. *Tcf7* reactivation upon stimulation withdrawal coincided with CD25 downregulation and CD62L upregulation, suggesting re-entry into a memory state (Fig 4D). To test whether *Tcf7* reactivation and reacquisition of memory potential also occurs *in vivo*, we transferred *Tcf7*-YFP low and high cells into naive recipient mice (Fig. 4A), and assayed their *Tcf7* expression at successive time points. Indeed, *Tcf7*-YFP low cells reactivated *Tcf7* expression progressively in the spleen and lymph nodes over 10 days with concomitant increases in CD62L and IL7Rα, indicating reacquisition of a memory phenotype (Fig. 4E-G; Fig. S4C-D).

**Figure 4:**
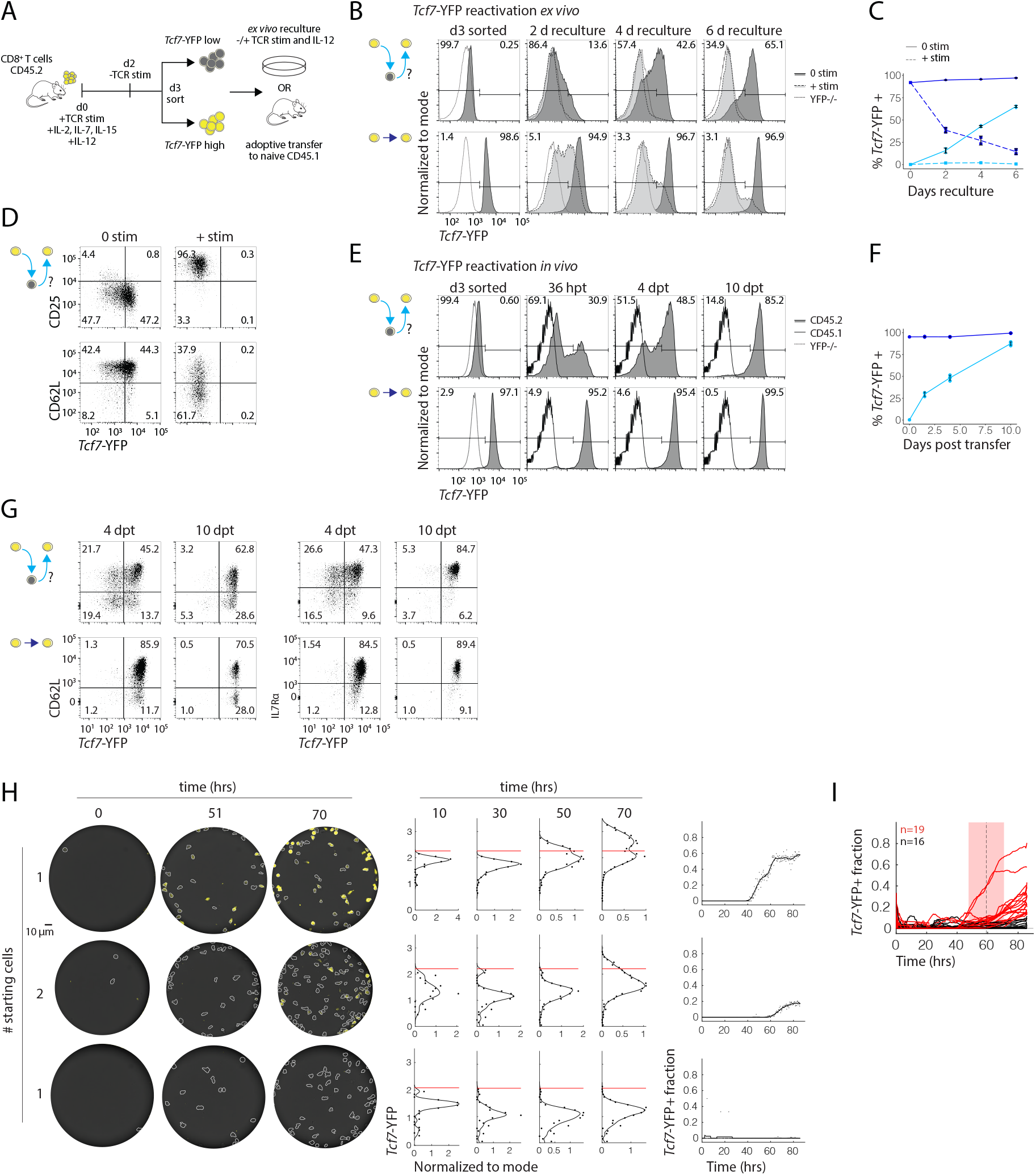
Effector cells reverse *Tcf7* silencing and regain memory potential upon stimulation withdrawal. (**A**) Naive cells from *Tcf7*-YFP mice were stimulated as indicated, sorted for *Tcf7*-YFP low and high populations after 3 days, and recultured either *ex vivo* (B-D, H-I) or adoptively transferred to naive recipients (E-G). Light and dark blue coloring throughout correspond to sorted YFP low and high populations, respectively. (**B-C**) *Tcf7*-YFP levels during reculture with or without αCD3/αCD28 and IL-12 (+/- stim) compared to non-fluorescent controls. **(D)**CD25, CD62L, and *Tcf7*-YFP expression in *Tcf7-low* sorted cells recultured +/- stimulation. (**E-F**) *Tcf7*-YFP levels over time in CD45.2^+^CD45.1^-^CD8^+^ cells harvested from the spleen after sort and adoptive transfer to naive recipients. (**G**) CD62L and IL7Rɑ expression in cells from E, F after *in vivo* transfer. (**H**) Representative microwells of sorted *Tcf7-low* cells recultured without stimulation: snap shots (left), top and bottom wells represent single clones; corresponding histograms (middle) with binned cell data for each time point, with YFP +/- gate drawn at 2 standard deviations above the mean YFP intensity from the first 25 hrs; corresponding YFP^+^ fractions over time (right). (**I**) YFP^+^ fraction for all wells overlaid. Mean activation time = 59.1 hr. [C, F] Mean ± s.d. [B-D] Data are from a single experiment representative of 1 and 3 independent experiments for +stim and 0 stim, respectively. [E-G] Data are from a single experiment with n=3-4 biological replicates.

We next used clonal live imaging of sorted *Tcf7*-YFP low cells confined in microwells to test if *Tcf7* reactivation is heterogeneous within individual effector clones, as would be expected if reactivation occurs via reversal of stochastic *cis*-epigenetic silencing (Fig. 3). Consistent with reactivation observed from bulk starting populations, individual *Tcf7* silenced cells gave rise to *Tcf7* high cells (Fig 4H-I; Supplementary Movies 2 and 3; Supplementary Table 5). Similar to the initial *Tcf7* silencing event, reactivation was heterogeneous within clones. Reactivation occurred only after multiple divisions, which may reflect the need for cell division for permissive chromatin state changes. Overall, these results indicate that cells that have silenced *Tcf7* and relinquished memory potential can reverse this decision later, after resolution of an immune challenge.

### *Tcf7* high cells formed through early and late decisions acquire a common memory program

Our results show that memory cells can arise through two pathways: a “naive to memory” (NM) pathway, whereby some cells maintain *Tcf7* expression during initial antigen stimulation, and a “naive to effector to memory” (NEM) pathway, by which cells that have silenced *Tcf7* and entered an effector state can turn expression back on after stimulation removal. To determine whether *Tcf7* high cells emerging through these two pathways both have genomic and functional memory programs, we subjected them to transcriptomic, epigenomic, and cytokine secretion analysis, alongside control *in vivo* naive (CD44^-^CD62L^+^), memory (CD44^+^CD62L^+^), and *ex vivo* generated effector cells (Fig. 5).

**Figure 5:**
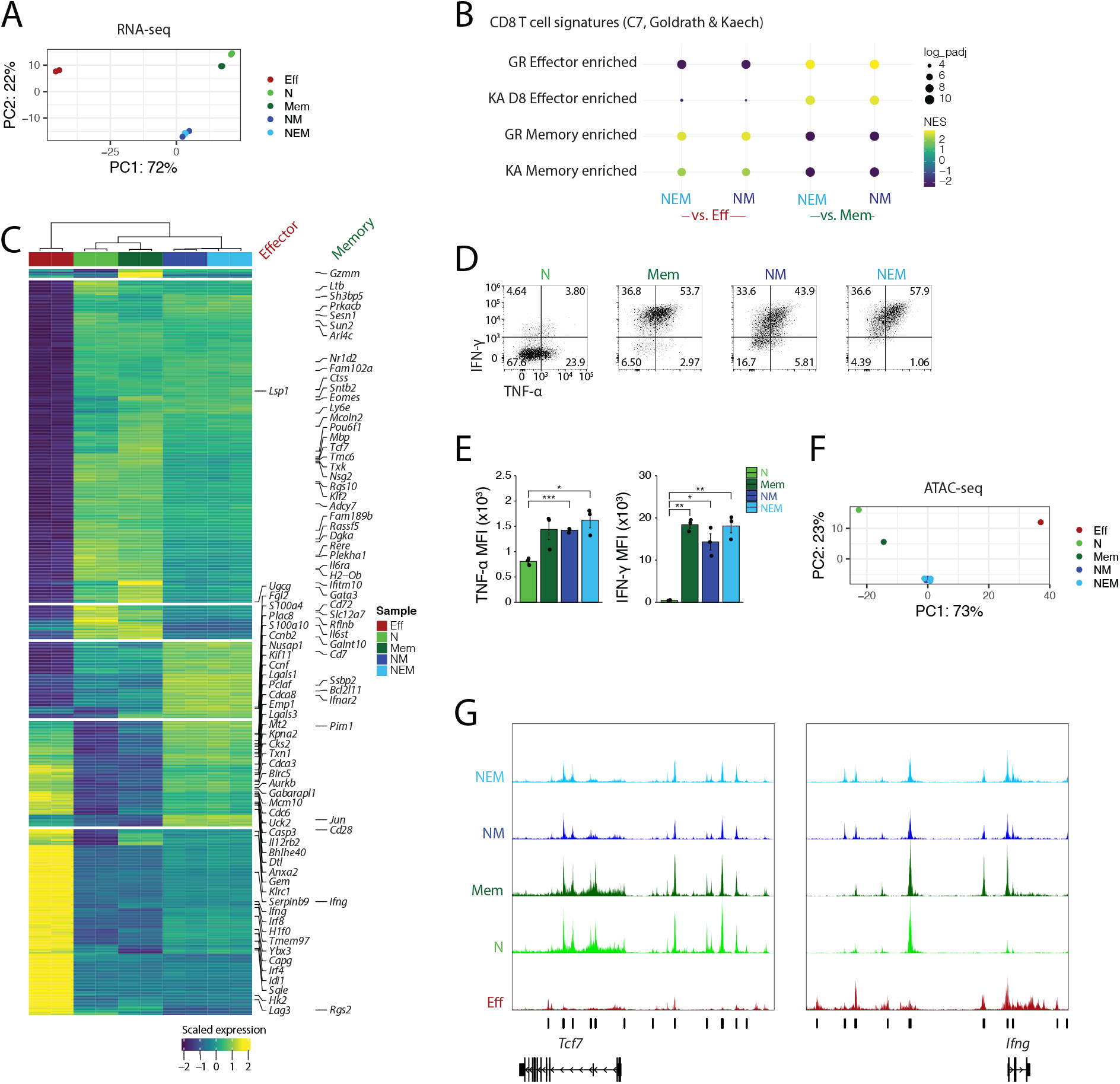
*Tcf7* high cells emerging from different routes acquire memory programming and functions. *Tcf7*-YFP low and high cells were sorted after 2 days of stimulation followed by one day of rest, recultured without TCR stimulation or IL-12 for an additional 6 days *ex vivo*,and then sorted for high *Tcf7*-YFP expression and subjected to genomic and functional analyses. (**A**) PCA of RNA-seq profiles (top 500 DE genes) for recultured cells compared to day 3 effector (Eff) and day 0 naive (CD44^-^CD62L^+^, N) and memory (CD44^+^CD62L^+^, Mem) controls. NM and NEM cells were sorted as YFP-high and YFP-low on day 3, respectively. (**B**) GSEA of gene signatures from MSigDB (C7, collections deposited by Goldrath (GR) and Kaech (KA) comparing *ex vivo* recultured populations to Eff and Mem controls. (**C**) Heatmap displaying top 500 DE genes (lfc ≥ 2, Bonferroni-adjusted p value <0.05) between recultured populations and Eff, N, and Mem controls. Scale bar indicates row z-scores of regularized log transformed count data. Memory and effector associated genes from MSigDB Goldrath and Kaech collections are highlighted. (**D-E**) Cytokine secretion of recultured cells compared to N and Mem controls after PMA/Ionomcyin restimulation. (**F**) PCA of ATAC-seq counts of top 500 differentially accessible peaks between recultured cells and controls. (**G**) ATAC-seq read coverage tracks; vertical bars annotate differentially accessible peaks between recultured cells and controls. [A-C] n = 2 biological replicates for each sample. [D-E] Mean ± s.d. Statistical significance was calculated with an unpaired two-tailed t test performed between groups. *p<0.5, **p<0.01, ***p<0.001. Data are n=3 biological replicates from a single experiment. [F-G] n = 1 biological replicate for Eff, N, Mem, n = 2 for NM, n = 3 for NEM.

Remarkably, NM and NEM cells showed similar memory characteristics, despite different *Tcf7* regulatory history. They were both more similar to naive and memory *in vivo* controls compared to *ex vivo* generated effector cells in their shared expression of memory-defining genes, though they also maintained some effector characteristics, in line with their recent stimulation (Fig. 5A-C; Fig. S5A). Similar to memory controls, NM and NEM cells demonstrated greater TNF-ɑ and IFN-γ secretion upon re-stimulation compared to naive cells (Fig. 5D-E). NM and NEM cells were most similar in global chromatin accessibility to memory controls (Fig. 5F; Fig. S5B-C). NEM cells recovered similar *Tcf7* accessibility to NM cells (Fig. 5G). At the *Ifng* locus, intermediate accessibility of NM and NEM cells between naive and effector controls suggests that both were poised for rapid recall response, and accessibility at other memory- and effector-associated loci support this conclusion (Fig. S5D).

While NM and NEM cells were largely similar, notable differences in tissue localization and gene expression suggest they may be primed for different functional memory properties *in vivo*. Transferred *Tcf7* high cells showed greater engraftment in secondary lymphoid organs than *Tcf7* low sorted cells, suggesting different homing capabilities (Fig. S5E). NEM cells also had higher expression and accessibility of some effector-associated genes compared to NM, possibly indicative of enhanced effector capabilities or an effector memory state^37,38^ (Fig. S5F-G). Overall, both NM and NEM decision strategies give rise to cells with genomic and functional characteristics of memory, suggesting that memory formation may proceed through a flexible decision-making strategy, allowing both for memory and effector divergence during the initial immune challenge and for effector reacquisition of memory potential after the challenge is resolved.

### Multiple paths to memory enable robust encoding of pathogen experience through memory population size

Flexibility in memory decision-making may have functional benefits, and may in particular allow for scaling of memory population sizes with immune response magnitudes. To test this idea, we used mathematical modeling to evaluate different T cell decision-making strategies in their memory outcomes in response to pathogens of different virulence, modeled as having different rates of replication (see Mathematical Appendix). In our first model, we consider the flexible strategy we observed (Fig. 6A). Here, naive T cells (*T_n_*) initially transition to a *Tcf7-*expressing memory-competent state (MC, *T_m_*) that divides upon exposure to pathogen (*v*), but stops dividing and persists upon pathogen clearance. These cells can either maintain memory competence upon continuing stimulation, or transition to *Tcf7-silent* effector state (*T_e_*), where they control pathogen growth, but are short-lived. Based on our findings (Fig. 3), this transition to an effector state is stochastic, with a probabilistic rate that increases with pathogen. Effector cells can reverse *Tcf7* silencing and re-enter the memory-competent state in the absence of pathogen, as we observe (Fig. 4).

**Figure 6:**
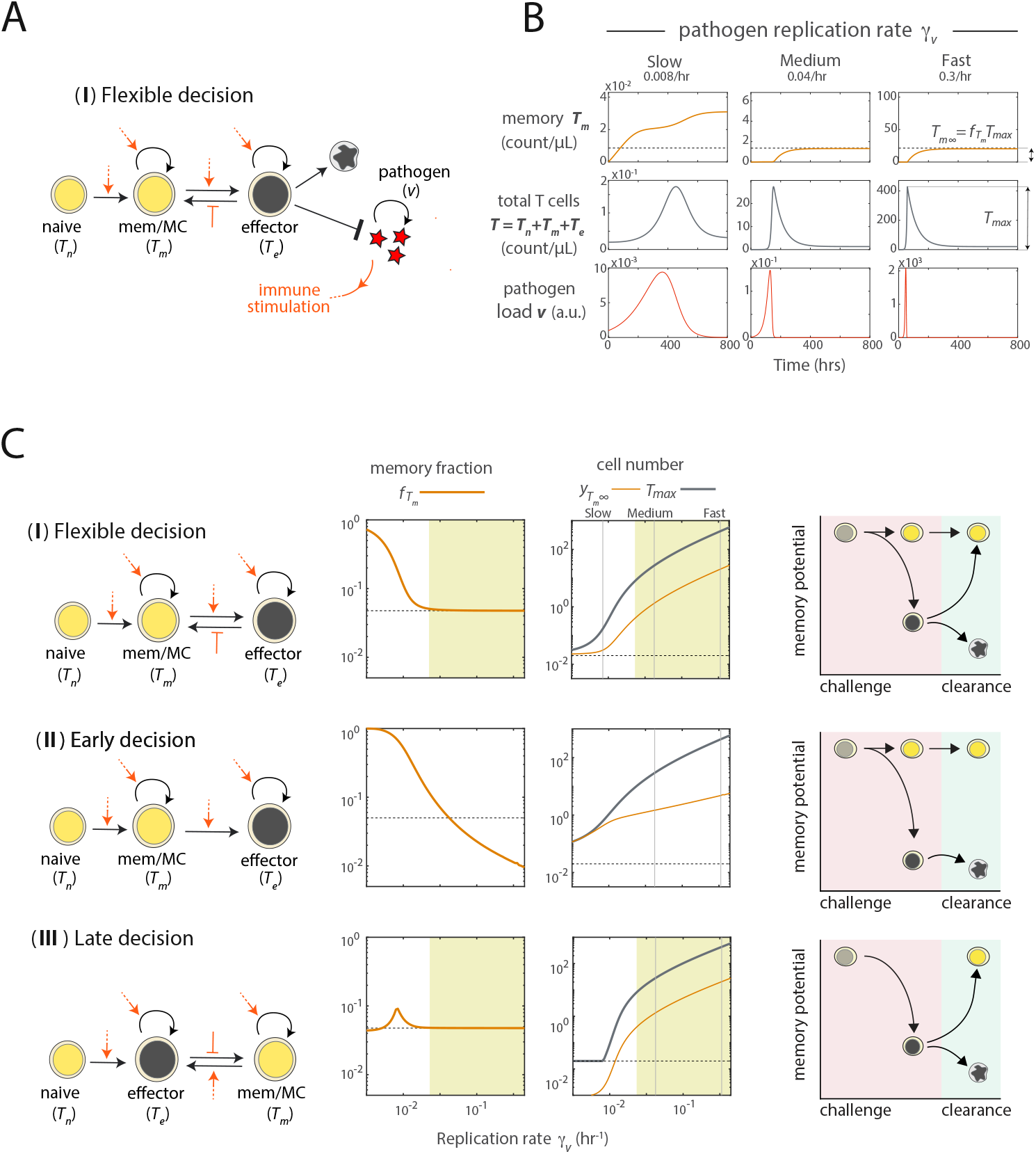
Flexible decision making enables quantitative encoding of pathogen experience during T cell memory formation. (**A**) Model incorporates pathogen proliferation, T cell memory decision making through reversible epigenetic switching. Orange arrows indicate modulation of T cell state transitions by pathogen load. (**B**) Time traces show memory T cell levels (top), total T cell levels (middle) and pathogen load (bottom), for different rates of pathogen replication (left to right). Dotted line shows the number of memory T cells formed in the case when this number is a defined fraction of the peak total T cell number, f_Tm_. (**C**) Distinct strategies for memory decision making: flexible (top), early (middle) or late (bottom); the fraction of T cells at the response peak that become memory cells f_Tm_; the peak cell number (black) and memory cell number (orange), both plotted against pathogen replication rate γ_v_. The dotted line indicates the number of starting naive cells, and the yellow shading marks scalable memory.

Mathematical simulations of this flexible decision model recapitulate the canonical features of the T cell response to acute infection (Fig. 6B; Fig. S6A-B). T cells expand rapidly in response to pathogen, reaching a peak 4-8 days after infection onset that consists mostly of effector cells, followed by a contraction to a stable, lower level of memory-competent cells (*T_m_*). Consistent with known studies^1,39^, the quantity of memory cells is ~5% of the peak cell number.

In response to pathogens with varying replication rates, this flexible decision model allows memory cells to form robustly and scale linearly with peak cell expansion numbers. Increasing effector expansion with faster pathogen replication was accompanied by a proportional increase in memory cells, such that the memory fraction relative to the peak T cell number remained constant (Fig. 6B and 6C – top, yellow shading, γ_v_>0.02/hr). This relation is given by:

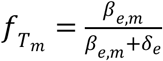

where *β_e,m_* is the maximum effector to memory conversion rate and *δ_e_* is the effector death rate. This scaling breaks down when pathogen replication is slow (γ_v_<0.02/hr): reduced antigen encounter decreases the probability of the early effector cell decision, such that the number of memory cells generated converges to the starting naive cell number rather than increasing with pathogen replication rate. This ensures a baseline level of memory amid weak challenges that do not elicit a full effector response^3^.

To ask whether flexibility is necessary for scalable memory encoding, we analyzed two alternative decision models, where memory decisions are made at only one juncture. The early decision model, where naive cells irreversibly commit to the *Tcf7*-silent effector state, generated robust memory upon challenge with slow-dividing pathogens but cannot reproduce the linear scaling of the memory population to the peak population in response to faster-replicating pathogens (Fig. 6C, middle; Fig. S6C-F; see also Mathematical Appendix). Conversely, the late decision model, where naive cells transition obligatorily to the effector state and decide later whether to regain memory competence, generated constant memory fractions upon stronger challenges but attenuated memory populations in response to weaker challenges (Fig. 6C, bottom; Fig. S6G-H). These analyses underscore the importance of flexibility in memory decision making for optimal long-term immunity against variable threats.

## Discussion

Our finding that reversible epigenetic silencing of *Tcf7* generates inherent flexibility in the T cell memory decision reconciles two prevailing models for memory development that have often been regarded as mutually opposed. While there is evidence that memory cells can form both directly from naive cells with little or no effector differentiation and from effector cells that dedifferentiate upon infection clearance^8,9^, no model has explained how both pathways can coexist. In this mechanism, stochastic control of *Tcf7* silencing enables early divergent memory and effector decision making, and its reversibility enables late effector dedifferentiation. Antigen and inflammatory signals tune the decision-making probabilities at both junctures (Fig. 2–4) and would thereby influence which pathway would predominate across challenges that differ in signal duration and intensity^40^. This study, together with others^33^, implicate stochastic epigenetic switches as drivers of cellular diversification in the immune system. Through regulatory events that initiate over timescales spanning cell generations, these switches allow multiple cell populations to emerge in defined numbers without strict spatially-organized cues^41^, facilitating division of labor for optimal pathogen defense.

Our modeling results lay the groundwork for understanding how the adaptive immune system can encode information about the nature and severity of a pathogen in its memory cell population (Fig. 6). In future work, it will be interesting to determine whether other pathogen features, such as antigenicity or latency, may also be encoded quantitatively. Our findings that memory cells emerging from different decision points may differ in their functional properties (Fig. S5E-G) raise the possibility that flexible decision making could underlie qualitative encoding of pathogen information through the generation of heterogeneous memory subsets^37,38^. In future work, it will be interesting to investigate the extent to which each decision pathway is utilized under various threats *in vivo* and whether cells emerging from different pathways are functionally heterogeneous^42^.

Overall, our study highlights the utility of plasticity in cell fate decision making in biological systems. From a strategic standpoint, flexibility enables decision makers to change their minds with new information, allowing them to mount optimal responses amid uncertain circumstances^12^. For immune cells responding to a pathogen, an ability to reassess prior decisions, as opposed to making early commitments, may enable bet-hedging and greater responsiveness as an immune challenge evolves. Observed plasticity in mammalian stem cell fate decision making^43,44^ may similarly allow the body to rapidly adapt its regenerative output to changing physiological needs^45^. A fuller consideration of flexibility in cellular decision making, along with its mechanisms and roles, will shed light into design principles of these systems and provide valuable insight for harnessing cells as environmentally-responsive therapeutic agents.

## Acknowledgments

We thank members of the Kueh, Nourmohammad, and Shendure labs, as well as Philip Greenberg, Jinfang Zhu, and Douglas Fowler for stimulating discussions and feedback, and William Noble for advice on statistical analysis methods. We also thank Sam Nguyen for help with brightfield image cell segmentation. This study was funded by an NIH/NIBIB Trailblazer Award (R21EB027327, to H.Y.K.) an NIH/NHGRI R01 grant (R01HG010632-01, to J.S.), NSF Graduate Research Fellowships (K.A., E.C.C.), NSF CAREER award (grant No: 2045054, to A.N.), and startup funds from the Bioengineering Department at the University of Washington (H.Y.K.), the Department of Physics at the University of Washington (A.N.), and the Rockefeller University (J.C.). J.S. is an investigator of the Howard Hughes Medical Institute.

## Author Contributions

K.A. and H.Y.K. conceived the study and designed the experiments. E.C.C., J.S. and J.C. contributed to experimental design. K.A. and E.C.C. performed the experiments and analyzed the data. J.C. performed the scRNA-seq experiments, and R.D. performed bulk RNAseq and ATACseq. W.Y. performed initial analysis on the scRNA-seq experiments. J.F. and A.L.W performed analysis on imaging data. K.K.H.N. helped set up the *ex vivo* T cell activation system. H.Y.K. developed the mathematical models and O.U., A.N., and H.Y.K. analyzed the mathematical models. A.B. provided the *Tcf7*-YFP reporter mice and guidance. K.A. and H.Y.K wrote the manuscript. E.C.C. contributed to the writing of the manuscript.

## Data availability

The time-resolved single-cell RNA-seq and bulk RNA-seq and ATAC-seq data generated for this study will be deposited in the Gene Expression Omnibus. All other data will be made available upon reasonable request.

## Code availability

Scripts for processing sequencing and imaging data are written in Python, R, and Matlab and will be made available at https://github.com/KuehLabUW and https://github.com/kathleenabadie.

## Methods

### Mice

*Tcf7-*YFP mice have been described^16^. We note that a small number of experiments utilized mice harboring an additional non-perturbing *Tbx21-CFP* BAC transgene reporter allele^46^, though this reporter was not further analyzed for this study. All mice used in experiments were heterozygous for the *Tcf7-*YFP reporter except where specified. WT C57BL/6 mice (Jackson Laboratory) were utilized as reporter negative controls, where applicable. Both male and female mice were used for *ex vivo* experiments, aged 8 to 12 weeks. Female CD45.1 mice, 8-12 weeks of age, were purchased from the Jackson Laboratory for use as recipients for adoptive transfer experiments. For donors for adoptive transfer experiments, homozygous *Tcf7*-YFP mice were crossed with an LCMV specific TCR transgenic strain^47^ (P14) (Jackson Laboratory) and heterozygous offspring were used. P14 homozygous mice without *Tcf7*-YFP were utilized as non-fluorescent controls for sort gate setting. All mice were used in accordance with Institutional Animal Care and Use Committee guidelines for the University of Washington.

### Naive T cell extraction

Spleens were harvested from mice, massaged between rough glass slides to generate a single-cell suspension, and filtered through 40 μm nylon mesh into HBH (HBSS, 10 mM HEPES, 0.5% BSA, pH 7.4). Cells were spun down for 5 min at 300g, resuspended in 3 ml red blood cell (RBC) lysis buffer (150 mM NH4Cl, 10 mM NaHCO3, 1 mM EDTA) for 3-5 min, and quenched with HBH. Cells were spun down for 5 min at 300g and resuspended in HBH with 2.4G2 blocking solution and incubated for 30 min on ice. Cells were counted, spun down again, and then enriched for CD8 T cells using a CD8a^+^ T Cell Isolation Kit, mouse (Miltenyi, #130-104-075), with the volume and amount of antibodies and microbeads used scaled down to 70% of that specified by the manufacturer. One LS column was used per spleen (Miltenyi, # 130-042-401). To obtain a pure population of naive CD8 T cells, the cell suspension was stained with anti-CD8 (PerCP/Cyanine5.5, eBioscience, # 45-0081-82 or Biolegend, #100734), anti-CD44 (APC or PE, Invitrogen, #17-0441-82, or #12-0331-82), and anti-CD62L (APC/eFluor780, Invitrogen, #47-0621-82) at 1:600 antibody to cell suspension volume ratio in 30×10^6^ cell/ml HBH with Fc block for 15-30 min on ice and then sorted with a BD FACS Aria III (BD Biosciences) with assistance from the University of Washington Pathology Flow Cytometry Core Facility. The naive population was gated as CD8^+^CD44^-^CD62L^+^*Tcf7*-YFP^+^. Memory cells were gated as CD8^+^CD44^+^CD62L^+^*Tcf7*-YFP^+^. The cells were sorted into HBH and kept on ice until plating.

### *Ex vivo* T cell differentiation

One day prior to T cell harvest and activation (day −1), plates were prepared by coating with anti-CD3e (Tonbo, #40-0031-U100), anti-CD28 (Tonbo, #40-0281-U100), RetroNectin (Takara, #T100B), and when specified, anti-CD11a (Biolegend, #101117). Unless otherwise specified, each well of a 96-well plate received 0.2 μg anti-CD3, 0.1 μg anti-CD28, 1 μg Retronectin, and (when specified) 1 μg anti-CD11a in 50 μL of PBS. For differentiation in larger wells, these amounts were scaled up by well surface area. Plates were sealed with parafilm and incubated at 4°C overnight. On day 0, plates were allowed to come to room temperature for at least 30 min and washed 2x with PBS. Purified cells were added to wells in T cell media [85% RPMI 1640 with L-glutamine, 10% Fetal Bovine Serum, Pen-Strep-Glutamine, 20 mM HEPES, 1 mM Sodium Pyruvate, 0.1 mM NEAA, 50 μM BME] with indicated cytokine concentrations, mixed, and spun down for 1 min at 150g to ensure initial contact for all cells with the coated plate surface. Cytokines added to the media were 100 U/mL IL-2 (PeproTech, # 200-02), 0.5 ng/mL IL-7 (PeproTech, # 200-07), 50 ng/mL IL-15 (PeproTech, # 210-15), and 1 ng/mL IL-12 (PeproTech, #210-12) unless otherwise specified. Where specified, IFN-β1 (Biolegend, #581302) was added at 1000 U/mL. The cell seeding concentration was 0.1 - 2.5 million cells / ml unless otherwise indicated. Cells were incubated at 37°C in 5% CO2 and split every two days by mixing, removing half of the well volume, and topping off the volume with TCM and respective cytokines. Where applicable, prior to seeding, cells were stained with 5 μM CellTrace Violet (CTV) (Invitrogen, #C34557) following the manufacturer’s instructions.

### Flow cytometry analysis

For timecourse analyses with cell surface protein staining, cells were spun down in round-bottom 96-well plates or 1.5 ml eppendorf tubes, resuspended in 2.4G2 blocking solution for 15-30 min on ice, stained with cell surface antibodies at 1:1200 (anti-CD8: PerCP-Cyanine5.5, eBioscience, # 45-0081-82, or Biolegend, #100734, anti-CD44: APC, Invitrogen, #17-0441-82, anti-CD62L: APC-e780, Invitrogen #47-0621-82, anti-CD25: APC, #17-0251-82), 1:400 (anti-CD45.1: APC, Biolegend, #110714), 1:200 (anti-CD45.2: PE/Dazzle594, Biolegend, #109846) or 1:100 (anti-CD127/IL7Rɑ: PE, Invitrogen, #12-1271-83) antibody to cell suspension volume ratio for an additional 15-30 min on ice, and spun down again for a final resuspension in HBH prior to acquisition. For samples that required intracellular protein staining, cells were fixed and permeabilized using Cytofix/Cytoperm Fixation and Permeabilization kit (BD #554714) according to manufacturer instructions and incubated with antibody for 30 min on ice. The TCF1 antibody (PE, BD Biosciences, #564271) and T-bet antibody (PE, Biolegend, #644809) were used at 1:50 and 1:200, respectively. For samples that required intracellular cytokine staining, cells were restimulated for 5 hr with PMA/Ionomycin (1x in 100 μL per sample Thermofisher, #00-4970-93) in round-bottom 96-well plates, with a protein transport inhibitor (1x Thermofisher, #00-4980-93) added after 1 hr. For cytokine secretion after sorting (for Naive, Mem, and NM/NEM) cells were stained with Zombie Near IR at a 1:1000 dilution in PBS following the manufacturer’s instructions (Biolegend, #423117). Cells were then fixed, permeabilized, and stained with antibodies for cytokine and other intracellular protein antibodies as described above. All cytokine antibodies were used at 1:100 dilution in 1x BD Perm/Wash buffer (anti-IFN-γ (APC/Cyanine7 or PE, Biolegend, #505849, #505808) and anti-TNF-ɑ (BV711 Biolegend, #506349). Data were acquired using an Attune Nxt Flow Cytometer (ThermoFisher Scientific) and analyzed using FlowJo (BD) software.

### Sample processing for sci-fate-seq

Naive CD8 T cells were activated *ex vivo*, as described. For this experiment, media was supplemented with 100 U/mL IL-2, 0.5 ng/mL IL-7, 50 ng/mL IL-15, and 0.05 ng/mL IL-12. The moderate level of IL-12, 0.05 ng/ml, was chosen for this experiment to produce a relatively even representation of *Tcf7* high and low cells (see Fig. 1C). At days 1, 2, and 4 of activation, two subsequent sci-fate time points were taken as follows: cells were mixed and split into two wells, which had been coated with anti-CD3 and anti-CD28 at day −1 and remained in the incubator with TCM; 4sU was added to one well for a final concentration of 200 μM, and that well was harvested 2 hr later. At that time, 4sU was similarly added to the second well, and that well was harvested 2 hr later. After each 4sU addition, cells were mixed and spun down at 150g for 1 min. Harvested cells were prepared for sci-RNA-seq as described for the sci-fate protocol ^22^. Briefly, cells were fixed with ice-cold 4% PFA for 15 min, washed and flash frozen with PBSR [PBS, pH 7.4, 0.2 mg/ml bovine serum albumin (Fisher), 1% SuperRnaseIn (Thermofisher) and 10 mM dithiothreitol (DTT)]. PFA-fixed cells were thawed, washed, and treated with iodoacetamide (IAA) to attach a carboxyamidomethyl group to 4sU. Following these steps, a single-cell RNA sequencing library was prepared using the sci-RNA-seq protocol^48,49^. The library was sequenced on the Illumina Next-seq system.

### Computational analysis for sci-fate-seq

#### Read alignment, downstream processing, and TF module construction

Read alignment and downstream processing, linking of TFs to regulated genes, and construction of TF modules was performed as described in Cao et al., 2020, with minor modifications. Briefly, for each gene, across all cells, the correlation between mRNA levels of each expressed TF and that gene was computed using LASSO (least absolute shrinkage and selection operator) regression. We sought to comprehensively define gene programs with distinct dynamics by doing this correlation separately both using only newly synthesized transcript levels for potential target genes and using overall transcript levels, expecting that target genes with more stable transcripts would be more readily identified using newly synthesized transcripts, while less abundant, more lowly detected target genes would be more readily identified in the overall transcriptome. After filtering out the resultant covariance links with a correlation coefficient less than 0.03, we identified 2,117 putative TF - target gene covariance links using newly synthesized transcriptome levels and 9,927 using overall transcriptome levels, resulting in a total of 10,405 unique links after aggregation. These were further filtered to retain only links supported by ChiP-seq binding, motif enrichment, or predicted enhancer binding^24^, resulting in 1065 links between 51 TFs and 632 genes. Of these 1065 links, 147 were identified using the newly synthesized transcriptome levels, 649 were identified using the overall transcriptome levels, and 269 were identified by both. To calculate TF activity scores in each cell, newly synthesized unique molecular identifier (UMI) counts for all linked target genes were scaled by library size, log transformed, aggregated, and normalized. The absolute correlation coefficient was computed between all TF pairs with respect to their activity across all cells. Pairwise correlations were hierarchically clustered using the ward D2 method to identify TF modules, with the reasoning that co-regulatory TFs must be simultaneously active within the same cell.

#### Cell ordering, clustering, and differential gene expression analysis between clusters

We initially attempted to resolve T cell differentiation states by performing dimensionality reduction with Uniform Manifold Approximation and Projection (UMAP) on whole or new transcriptomes using all detected genes. This analysis largely separated cells by the time point at which they were sampled (Fig. S2B), as previously observed^50,51^, likely a consequence of the host of other temporal changes occurring during activation apart from differentiation, such as cell cycle control and metabolic programming. To characterize T cell differentiation dynamics apart from other regulatory processes, cells were represented in UMAP space using newly synthesized reads for all genes within the T cell differentiation TF module with monocle3 (v.0.2.3.0) (reduction_method = ‘UMAP’, umap.n_neighbors = 15L, umap.min_dist = 0.001) ^52^ using the function align_cds^53^ to remove effects of cell cycle phase (preprocess_method = ‘PCA’, alignment_group = ‘Phase’). The resultant UMAP was clustered using density peak clustering^54^, which resulted in 5 main clusters (Fig. 2C, U and E2(A) combined, E1(A), E1(B), E2(B), and MP(A) and MP(B) combined). To further resolve observed variable T cell differentiation marker expression within two of these clusters, *k-*means clustering was used to further divide U and E2(A) into separate states and MP(A) and MP(B) into separate states (*k* = 2 and 2.5, respectively). Cells in different cell cycle phases were relatively evenly distributed across this UMAP, with S phase representation highest in E1(A) (Fig. S2D). Differential gene expression testing was performed between clusters using the monocle3 fit_models function.

#### RNA velocity analysis

RNA velocity analysis and visualization of velocity streamlines was performed using Dynamo (v.0.95.2.dev)^26^ using expression matrices from the full and new transcriptome. The dataset was subsetted to include only the T cell differentiation module genes prior to analysis, but the resultant streamlines were similar when the analysis was performed with all genes. The streamline results were also similar when scVelo (v.0.2.2)^55^ was used for velocity analysis (data not shown), with the full and new transcriptome used as the unspliced/spliced expression matrices, indicating that the streamline results are consistent between multiple analysis methods. The scVelo results were also similar with or without subsetting to include only the T cell differentiation module genes.

#### Trajectory analysis

Cells in each putative trajectory (E1, E2, MP) were ordered in pseudotime based on the point position on the principal curve estimated using the princurve package^56^. To align the precursor cells between trajectories, cells in the undifferentiated (U) cluster were set to pseudotime = 0. To identify genes that distinguish the trajectories, differentially expressed genes were identified using the monocle3 fit_models function with the model formula as the trajectory and pseudotime terms. Only resulting DEG associated with the trajectory term were selected.

### Time-lapse imaging

Long-term time-lapse imaging of cultured cells, both to track *Tcf7* regulation during initial activation in naive cells and to track *Tcf7* reactivation in sorted *Tcf7*-low cells, was performed as previously described with some modifications^57,58^. Images were acquired with an inverted widefield fluorescence microscope (Leica DMi8) fit with an incubator to maintain a constant humidified environment at 37°C and 5% CO2, using a 40X dry objective. For imaging of the initial 4 days of activation (Fig. 3), cells were seeded at low density (2-5k c/well) in wells of a 96-well glass bottom plate (Mattek) coated with anti-CD3, anti-CD28, anti-CD11a, and RetroNectin, as described above. For *Tcf7* reactivation imaging experiments (Fig. 4), *Tcf7*-low cells were sorted on day 3 after 2 days of initial culture with anti-CD3 and anti-CD28 in media with IL-2, IL-7, IL-15, and IL-12 and one additional day of culture with anti-CD3 and anti-CD28 removed. These cells were seeded onto PDMS micromesh (250 μm hole diameter, Microsurfaces) mounted on top of a 24-well glass bottom plate (Mattek) to enable clonal tracking, as seeded cells show considerably enhanced motility in the absence of TCR stimulation. To prepare the micromesh for imaging, the surface was first coated with BSA while mounted on top of a 24-well plate overnight at 4°C and then transferred to a new glass well and coated with anti-CD11a and RetroNectin for improved adhesion but without anti-CD3 and anti-CD28. For reactivation experiments, cells were cultured in TCM with IL-2, IL-7, and IL-15, but without IL-12.

To determine if the experimental conditions required for imaging affect differentiation, we systematically compared expression of CD44, CD62L, and *Tcf7*-YFP in cells activated on glass or tissue culture plates, at high or low seeding density, and with or without presence of anti-CD11a (Fig. S3L). CD44 levels were comparable across all conditions, confirming that all cells activated in all conditions. In tissue culture plates, CD62L and *Tcf7*-YFP levels were also comparable, though the *Tcf7*-YFP levels were slightly reduced at lower cell density, particularly in the condition without IL-12, consistent with previous findings that memory differentiation occurs less efficiently at lower cell densities^59^. On glass plates, the fraction of CD62L low cells was increased compared to on tissue culture plates. *Tcf7* levels were similarly low for the condition with IL-12, but the combination of low seeding density and presence of anti-CD11a on the glass plate resulted in a lower *Tcf7* distribution in the no IL-12 condition than was otherwise observed. This analysis shows that the specific conditions used for imaging do not affect overall differentiation trends but may underestimate the differences in differentiation between conditions with and without IL-12.

### Computational analysis for time-lapse imaging

#### Image segmentation and tracking

Image pre-processing, cell segmentation, and tracking was performed in MATLAB (Mathworks, Natick, MA) using the ictrack movie analysis pipeline we described previously^58,60^ (Fig. S3A-B), modified to enable segmentation of cells from brightfield movies. Importantly, to segment cells without additional fluorescent labels besides *Tcf7*-YFP, we first trained a convolutional neural network (CNN) with a U-net architecture^61^ to predict fluorescence images of whole cells from brightfield images, using images of cell-trace violet labeled T cells as a training data set^27^. We trained separate CNNs for the images acquired in 96-well plates (Fig. 3) and in microwells (Fig. 4), as predictions are optimal when images for training and prediction have similar features. For each training dataset, hundreds of images of CTV-stained cells were acquired at multiple timepoints during the process of interest (e.g. initial T cell activation or culture after stimulation removal). Using the trained CNN, we then generated predicted whole-cell fluorescence images from acquired brightfield movies, which were used for cell segmentation (Fig. S3B, 1.). Briefly, in the ictrack analysis pipeline, images underwent (1) correction by subtraction of uneven background signal stemming from the bottom of the glass plate or the side of the PDMS microwells (2) Gaussian blur followed by pixel value saturation to fix uneven signal intensity within the nucleus of the cell and (3) Laplacian edge detection algorithm to identify the nucleus boundary. Non-cell objects were excluded via size and shape limit exclusions. To generate clonal lineage trees, cells were tracked automatically between adjacent movie frames using the Munkres assignment algorithm, and the resulting cell tracks were manually checked for errors and to annotate cell divisions (Fig. S3B, 2.).

#### Tcf7 *promoter state assignment and analysis*

To enable quantitative analysis of *Tcf7* promoter activity in clonal cell lineages, we assembled separate full tracks of total *Tcf7*-YFP fluorescence levels from the starting cell to each ending cell within a lineage tree, for all lineage trees analyzed (Fig. S3B, 3.). Fluorescence levels are halved at each cell division; thus, to ensure continuity in *Tcf7-*YFP levels in these tracks, we calculated for each parent-daughter cell pair an offset in *Tcf7-*YFP levels, that we added to the daughter cells and their progeny, as previously implemented^32^. These ‘continuized’ tracks were then smoothed using MATLAB medfilt1 (N=5) and smooth (span = 80 time points, equivalent to 20 hours, method = lowess), and their first derivatives with respect to time were calculated to generate single-cell tracks of *Tcf7* promoter activity for downstream HMM analysis (Fig. S3B, 4.).

Cell tracks were exported from MATLAB to R for downstream processing. *Tcf7* promoter states for each cell and time point were called from tracks of *Tcf7*-YFP level derivatives using Hidden Markov Model (HMM) modeling, implemented with the msm Package for R (v1.6.9)^62^. We initially tested four candidate HMM models with either three or four promoter states and variable constraints on the derivative ranges within each state (Fig. S3C-D). For each model, we constrained the mean and variance in *Tcf7* promoter activities of each state by fitting Gaussian distributions to the *Tcf7*-YFP derivatives at different time windows, to reflect our observations that cells are expected to be mostly in an inactive, active, or attenuated state at different times. We then compared the performance of these four models by calculating their log-likelihood and corresponding AIC (Akaike information criterion) scores. We also checked the quality of each model’s fit to the data by assessing whether residuals of the fit follow a Gaussian distribution^63^ (Fig. S3E). Based on this analysis, we chose a model in which cells transition between 4 states: off (initial), low active, high active, and off (Fig. 3B, Fig. S3F), and all start in the off-initial state at the beginning of the track. This four-state model performed favorably compared to other models, likely because it better accounts for the distinct distributions of promoter activity of silent and active cells at initial and later time points.

Using this four-state model, we assigned promoter activity states at each time point for each cell, removing potentially spurious transient promoter states by finding all promoter states lasting less than 8 hours and replacing them with the previously assigned promoter state. From these states, we then identified promoter silencing events as those involving a switch from active (high or low) to an inactive (off) state, and activation events as those involving a switch from inactive (off-initial or off) to active (high or low) states. We did not allow transitions back to the starting inactive (off-initial) state, as this state has a distinct *Tcf7* promoter activity distribution from the later silent state (off), likely reflecting the distinct noise characteristics of *Tcf7-*YFP levels at different stages after activation.

For analysis of *Tcf7* silencing between sister cells, we first assigned an ending cell state to all cells in the dataset, representing the final promoter state of the cell prior to division or the end of the cell track. Cells with a tracked duration of less than 3 hours and parents with ending cell state durations of less than 10 hours were also excluded, to ensure the analysis only includes sufficiently tracked cells and durable promoter states. We then collected all division events for which the parent cell was in an ON promoter state prior to division and asked whether the daughter cell tracks ended in an ON or OFF promoter state. We thus calculated the number of division events that lead to no (ON/ON), unequal (ON/OFF), or concordant (OFF/OFF) daughter silencing and then calculated the fractions of each category in the entire dataset and within each generation. To statistically analyze the degree of discordance in *Tcf7* silencing decisions between sister pairs by modifying Cohen’s kappa statistical test for inter-rater reliability as follows: division events were categorized as concordant (ON/ON or OFF/OFF) or discordant (ON/OFF) between sisters. The modified Cohen’s kappa coefficient, κ’, was calculated as the observed percentage of discordant events minus the percentage of discordant events expected by chance, divided by 1 minus the percentage of discordant events expected by chance^64^ (Supplementary Table 4).

### Analysis of *Tcf7*-YFP negative fractions in homozygous and heterozygous reporter cells

For analysis in Fig. 3I-J and Fig. S3K, YFP distributions were exported from FlowJo as csvs, imported to Python, and represented as histograms. The positive and negative populations were fit simultaneously as two gaussian distributions using the scipy.optimize.least_squares function (scipy v1.5.2), and the gate between YFP positive and negative populations was identified as the intersection between the gaussian curves. The silent fraction was then calculated as the sum of the histogram below the gate divided by the sum of the entire histogram. Two-tailed unpaired t tests between homozygous and heterozygous YFP silent fractions were performed using scipy.stats.

### Sort and adoptive transfer or *ex vivo* reculture of activated cells

Cells were cultured *ex vivo* in the presence of anti-CD3/28 (+TCR stim) and IL-2, IL-7, IL-12, and IL-15 as described. On day 2, cells were transferred to a non-antibody-coated plate (-TCR stim) but kept in the same cytokine environment until day 3 for sorting. For adoptive transfer experiments only, CD8 T cells were activated directly after purifying with the Miltenyi CD8a^+^ T Cell Isolation Kit using 100% recommended reagent amounts, without further purifying naive starting cells by sorting, and RetroNectin was not used during anti-CD3/anti-CD28 stimulation. Prior to *ex vivo* activation, cells were stained with 2 or 5 μM CellTrace Violet (Invitrogen, #C34557). For *ex vivo* reculture experiments, cells were sorted from a single CellTrace peak representing cells that had undergone the same number of divisions over the 3 day culture period, to ensure YFP differences were due entirely to *Tcf7* regulation differences and not cell division differences. Cells were recultured with and without TCR stimulation and IL-12 (with IL-2, IL-7, and IL-15 maintained except where specified), as labeled in each figure, for an additional 6-10 days. For genomics experiments, effector controls (Eff) were activated with TCR stimulation and cytokines for 3 full days. For adoptive transfer, cells were sorted that had undergone at least 4 divisions. Cells were sorted on CellTrace Violet and *Tcf7*-YFP levels. The *Tcf7* low gate was set using wild type non-fluorescent control cells that were stimulated identically *ex vivo*. Using this negative gate, the top and bottom 20% of the YFP distribution were selected as *Tcf7* high and low. Sorted cells were resuspended in PBS and transferred retro-orbitally (1 million cells transferred per recipient) to naive CD45.1 mice. On days 1.5, 4, and 10 after adoptive transfer, mice were euthanized, and blood, spleens, and lymph nodes were collected for flow cytometry.

### Blood and lymph node processing

Blood was collected from euthanized mice by cardiac puncture. Red blood cells were lysed 2x for 5 minutes at room temperature using 1x RBC lysis buffer (described in naive T cell extraction), prior to proceeding with cell staining as described in Flow Cytometry Analysis. Inguinal lymph nodes were harvested, and massaged over a 40 μm cell strainer and resuspended for flow cytometry staining as described in Flow Cytometry Analysis.

### Sample processing for RNA-seq

Cells were centrifuged at 500g for 5 minutes, resuspended in 350 μL of Trizol (Ambion), mixed well, and frozen at −80°C for processing, starting from step 2 of the RNeasy micro kit (Qiagen, #74004) following the manufacturer’s instructions. After processing, RNA was resuspended in RNase free water, quantified using a NanoDrop 2000c (Thermo Scientific), and shipped on dry ice to Novogene Corporation Inc. (Sacramento, CA) for library preparation and sequencing.

### Computational analysis for RNA-seq

Raw FASTQ files from RNA-seq paired-end sequencing were aligned to the GRCm38/mm10 reference genome using Kallisto (v0.46.1)^65^, and the resultant transcript-level abundance estimates were imported to genes by cells matrices using tximport (v1.18.0) for downstream analysis. Transcripts with low counts (<10) were removed. Differentially expressed genes were identified with DESeq2 (v1.30.1)^66^. PCA plots were generated using the top 500 differentially expressed genes between NM and NEM samples and naive, memory, and effector controls. Significantly differentially expressed genes were also used for gene set enrichment analysis, performed with fgsea (v1.16.0)^67^ and using gene sets from the C7 immunologic or the H Hallmark gene-sets from Molecular Signatures Database deposited by Goldrath and Kaech.

### Sample processing for ATAC-seq

After sorting, cells were centrifuged at 500g for 5 minutes then supernatant was aspirated without disturbing the pellet. The pellets were resuspended in 100 μL of ATAC freezing buffer^68^ (50 mM Tris at pH 8.0, 25% glycerol, 5 mM Mg(OAc)2, 0.1 mM EDTA, 5 mM DTT, 1× protease inhibitor cocktail (Roche-noEDTA tablet), 1:2,500 superasin (Ambion)), flash frozen in liquid nitrogen and stored at −80°C. On the day of processing, samples were thawed, centrifuged at 4°C 500g for 5 minutes, and washed with 100 μL of cold 1X PBS. Cells were again centrifuged and resuspended in 100 μL Omni lysis buffer^69^ (RSB with 0.1% NP40, 0.1% Tween 20 and 0.01% Digitonin) and incubated on ice for 3 minutes, then quenched with 500 mL of RSB + 0.1% Tween 20. Nuclei were pelleted at 500g for 5 minutes at 4°C, resuspended in 100 μL cold PBS and counted. 50,000 nuclei were used per reaction, pelleted (500g for 5 min at 4°C), resuspended in tagmentation master mix^69^ (50 μL total: 25 μL 2X TD buffer, 16.5 μL 1x DPBS, 0.5 μL 1% Digitonin, 0.5 μL 10% Tween 20, 5 μL water, 2.5 μL Tn5 enzyme), and incubated at 55°C for 30 minutes. Samples were purified using DNA Clean and Concentrate-5 (Zymo Research) and eluted in EB buffer (10 mM Tris) for amplification of tagmented DNA. PCR amplifications were performed using Illumina indexed primers and NEBNext High-Fidelity 2X PCR Master Mix. SYBR green was added to each PCR reaction to monitor amplification before it reached saturation. Samples in this study were amplified between 11-15 cycles using recommended conditions^70^. Unpurified products were run on a 6% TBE gel for quality control. PCR product/library were purified using DNA Clean and Concentrate-5 (Zymo Research) then ran on a tapestation to visualize nucleosome distribution. The libraries were normalized to 2nM then pooled equimolar for sequencing. Pooled libraries were loaded onto a NextSeq 500 High150 cycle kit at 1.5 pM loading concentration with paired ends sequencing (read 1: 74 cycles, read 2: 74 cycles, index 1: 10 cycles, index 2: 10 cycles).

### Computational analysis for ATAC-seq

Raw ATAC-seq FASTQ files from paired-end sequencing were processed and aligned to the mm10 mouse genome using the PEPATAC (v0.10.3)^71^ pipeline, which uses bowtie2^72^ for alignment. Unmapped, unpaired, and mitochondrial reads were removed. Following alignment, peak calling, merging across all samples, and annotation was performed using HOMER (v4.10)^73^. Differentially accessible regions were identified using DESeq2. PCA plots were generated using the top 500 differentially accessible regions between recultured samples and naive, memory, and effector controls. Coverage tracks were generated from bigwig read alignment files using karyoploteR (v1.14.1).

### Statistical Analysis

All analyses and p or adjusted p value significance are listed with each figure caption. Statistics were performed in R using the rstatix package (v0.7.0) or Python using scipy (v1.5.2).

## Supplementary tables

**Table S1: Differential gene expression between all UMAP clusters in Fig. 2C.**These DEG results are displayed in heatmap in Fig. 2D.

**Table S2: Differential gene expression results for pairwise comparisons between relevant UMAP clusters in Fig. 2C.**These DEG results are displayed in volcano plots in Fig. 2E and Fig. S2F.

**Table S3: Differential gene expression between E1, E2, and MP trajectories.**These DEG results are displayed in Fig. 2G.

**Table S4: Discordance score calculation using modified Cohen’s kappa coefficient.**Results are displayed in Fig. 3H.

**Table S5: Analysis of *Tcf7*-YFP reactivation in microwells.**The number of microwells with a given starting number of cells and the number of microwells with this number of starting clones that gave rise to *Tcf7*-YFP reactivated cells is shown. Relevant to Fig. 4H-I.

**Table S6: Differentially expressed genes in each bulk RNA-seq cluster.**Relevant to Fig. 5C.

## Supplementary movies

**Movie S1: *Tcf7*-YFP silencing within a clonal lineage.**Relevant to Fig. 3A.

**Movie S2: *Tcf7-*YFP reactivation example 1.**Relevant to Fig. 4H-I.

**Movie S3: *Tcf7-*YFP reactivation example 2.**Relevant to Fig. 4H-I.

