## Supplementary Figures and Text for "Flexible and scalable control of T cell memory by a reversible epigenetic switch"

**This document includes the following supplementary materials**

Figs. S1 to S6

Mathematical Appendix

Supplementary references

#### Supplementary figures

Figure S1

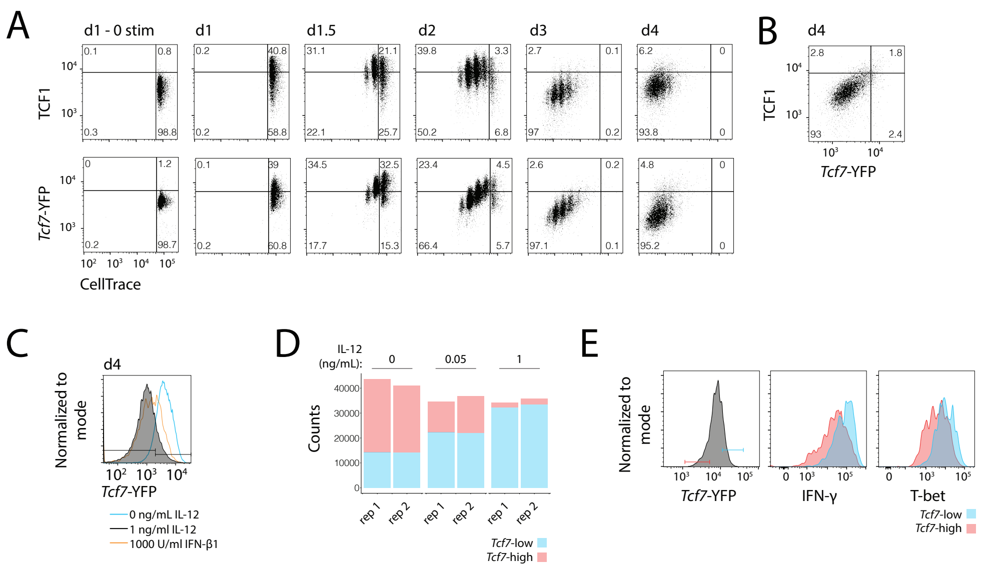


**Figure S1: Characterization of the *Tcf7*-YFP reporter during *ex vivo* CD8 T cell activation.** (**A-B**) TCF1 and *Tcf7*-YFP correlation in cells activated with 1 ng/ml IL-12. Note, *Tcf7*-YFP dynamic range is reduced in fixed and permeabilized samples due to leakage of fluorescent protein out of permeabilized cells. (**C**) *Tcf7* silencing in cells stimulated with IFN-β1 or IL-12. (**D**) Total *Tcf7*-YFP high and low cell counts for samples activated for 4 days with different IL-12 levels. (**E**) IFN-ɣ and T-bet levels in cells stimulated for 2 days with IL-12. Data are from a single experiment representative of [A-B] 2 independent experiments, [C] 1 experiment for IFN-β1 and at least 3 for others, [D] at least 3 independent experiments, [E] 1 experiment.

Figure S2

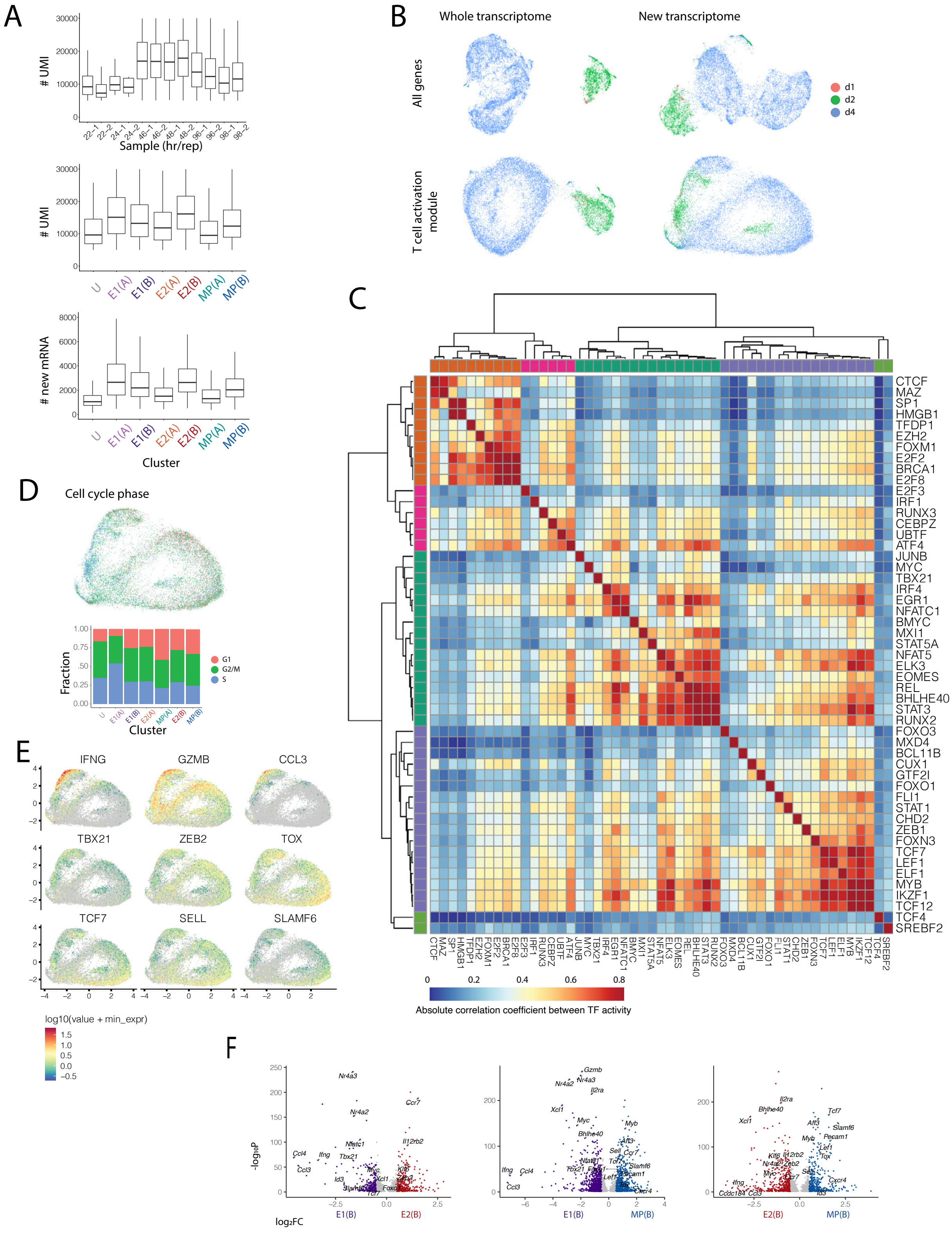
A
### UMI
Sample (hr/rep)
22-1
22-2
24-1
24-2
46-1
46-2
48-1
48-2
96-1
96-2
98-1
98-2
### UMI
### new mRNA
U
E1(A)
E1(B)
E2(A)
E2(B)
MP(A)
MP(B)
Cluster
B
Whole transcriptome
New transcriptome
All genes
T cell activation module
d1
d2
d4
C
CTCF
MAZ
SP1
HMGB1
TFDP1
EZH2
FOXM1
E2F2
BRCA1
E2F8
E2F3
IRF1
RUNX3
CEBPZ
UBTF
ATF4
JUNB
MYC
TBX21
IRF4
EGR1
NFATC1
BMYC
MXI1
STAT5A
NFAT5
ELK3
EOMES
REL
BHLHE40
STAT3
RUNX2
FOXO3
MXD4
BCL11B
CUX1
GTF2I
FOXO1
FLI1
STAT1
CHD2
ZEB1
FOXN3
TCF7
LEF1
ELF1
MYB
IKZF1
TCF12
TCF4
SREBF2
Absolute correlation coefficient between TF activity
D
Cell cycle phase
Fraction
G1
G2/M
S
Cluster
E
IFNG
GZMB
CCL3
TBX21
ZEB2
TOX
TCF7
SELL
SLAMF6
log10(value + min_expr)
F
-log10P
log2FC
E1(B)
E2(B)
MP(B)

**Figure S2: *Sci-fate* metrics and TF module analysis.** (**A**) Number of UMI per cell for each time point and replicate sample (top); number of UMI (middle) and newly synthesized reads (bottom) per cell for each cluster. (**B**) UMAP projections of cells using whole or new transcriptome and all genes or only T cell differentiation module genes. (**C**)Transcription factor module analysis as in Fig. 2B, enlarged to show direct TF correlations. (**D**) UMAP projection with cells colored by cell cycle phase[1] and fraction of cells in each phase for all clusters. (**E**) Expression of representative genes in UMAP space. **(F)** Differentially expressed genes between indicated clusters, using cells from all timepoints; log2FC > 0.5 and adj. p < 0.05.

Figure S3

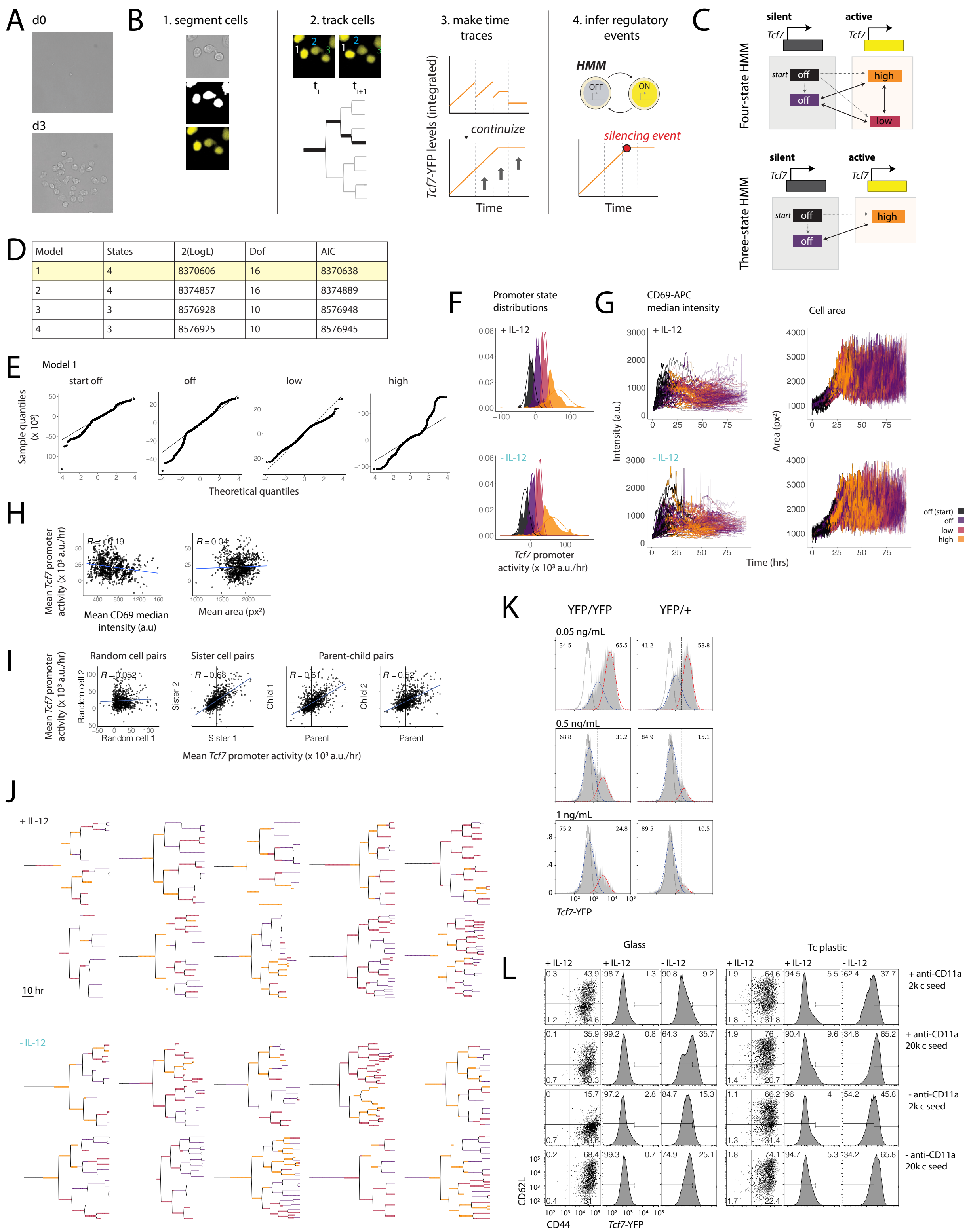


| Model | States | -2(LogL) | Dof | AIC |
|---|---|---|---|---|
| 1 | 4 | 8370606 | 16 | 8370638 |
| 2 | 4 | 8374857 | 16 | 8374889 |
| 3 | 3 | 8576928 | 10 | 8576948 |
| 4 | 3 | 8576925 | 10 | 8576945 |

**Figure S3: Quantitative live imaging reveals dynamics of epigenetic *Tcf7* silencing in clonal lineages.** (**A**) Representative images of a single clone adhered to the imaging plate at day 0 and day 3. (**B**) Quantitative image analysis procedure. (**C**) Four and three-state HMMs were compared and (**D**) evaluated by log-likelihood and AIC. Models 2 and 3 compared to models 1 and 4 have tighter constraints on the promoter activity range of the off states. (**E**) For each state in the selected HMM (model 1), quantiles of the theoretical gaussian distribution versus the quantiles of the observed residuals. The observed residuals would fall on the straight line in a perfect fit[2]. (**F**) *Tcf7*-YFP derivative, or promoter activity, of cells assigned to each HMM-derived promoter state. Theoretical distributions for each are overlaid as lines. (**G**) Median CD69 intensity and cell mask area for all overlaid tracks, colored by promoter state. (**H**) Average *Tcf7* promoter activity versus average CD69 median intensity and area for each cell trace, where a cell trace is an ending cell tracked from its naive progenitor. (**I**) Promoter activity averaged over entire cell cycle for related cells. (**J**) Additional lineage trees, as in Fig. 3A. (**K**) YFP distributions for homozygous and heterozygous reporters cultured for 5 days. (**L**) CD62L x CD44 and *Tcf7*-YFP distributions for varied activation conditions after 4 days of stimulation. [K] Data are from a single experiment representative of 2 independent experiments. [L] Data are representative of one experiment.

Figure S4

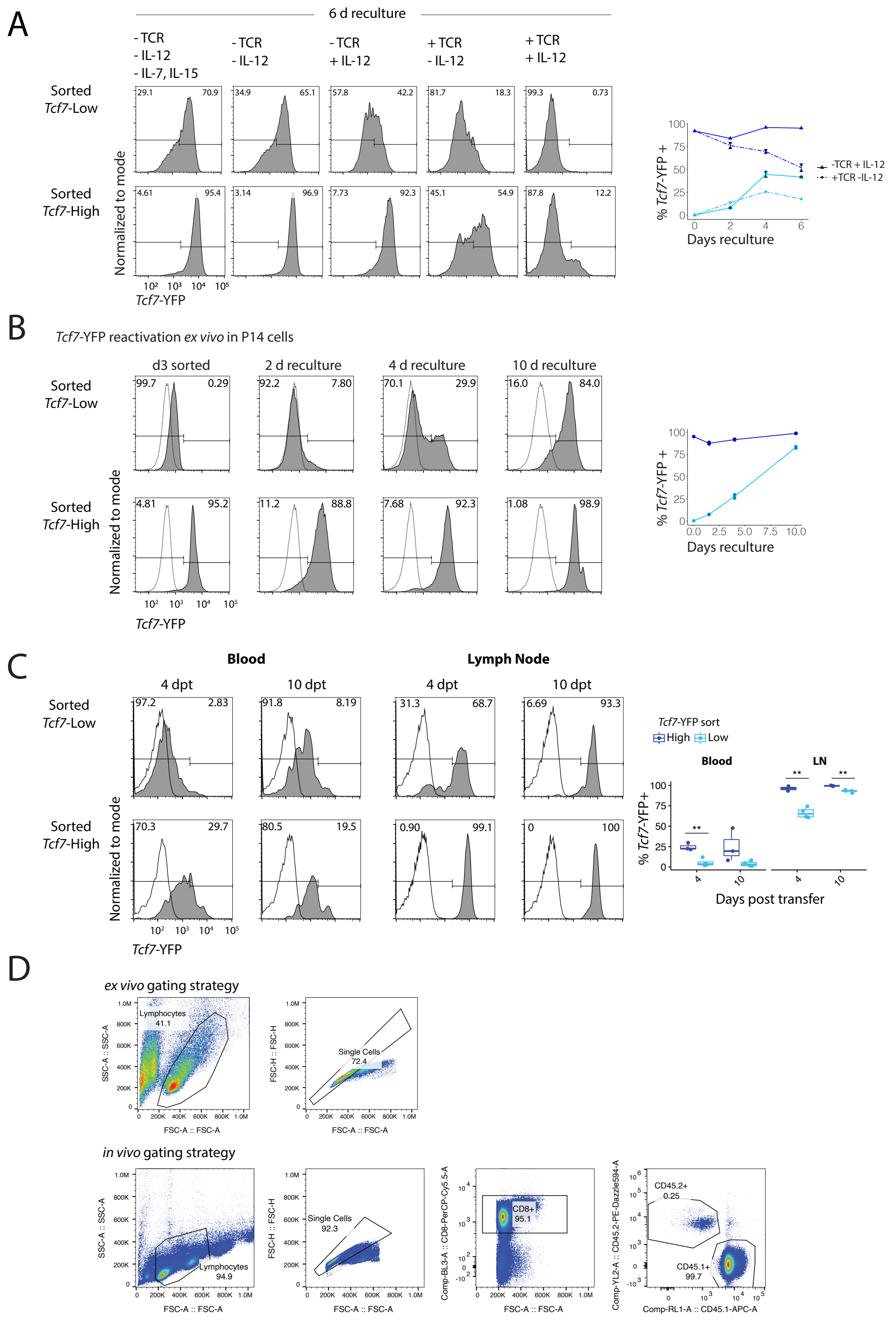
A
6 d reculture
- TCR
- IL-12
- IL-7, IL-15
+ TCR
+ IL-12
Sorted Tcf7-Low
Sorted Tcf7-High
Normalized to mode
Tcf7-YFP
% Tcf7-YFP +
Days reculture
-TCR + IL-12
+TCR -IL-12
B
Tcf7-YFP reactivation ex vivo in P14 cells
d3 sorted
2 d reculture
4 d reculture
10 d reculture
C
Blood
Lymph Node
4 dpt
10 dpt
Tcf7-YFP sort
High
Low
LN
% Tcf7-YFP+
Days post transfer
D
ex vivo gating strategy
Lymphocytes
41.1
Single Cells
72.4
in vivo gating strategy
Single Cells
92.3
Lymphocytes
94.9
CD8+
95.1
CD45.2+
0.25
CD45.1+
99.7
SSC-A :: SSC-A
FSC-A :: FSC-A
FSC-H :: FSC-H
Comp-BL3-A :: CD8-PerCP-Cy5.5-A
Comp-YL2-A :: CD45.2-PE-Dazzle594-A
Comp-RL1-A :: CD45.1-APC-A

**Figure S4: *Tcf7* reactivation upon signal withdrawal.** (**A**) Naive *Tcf7*-YFP low and high cells were sorted after 2 days of TCR stimulation with IL-12 and one day with removal of TCR stimulation. Sorted populations were recultured *ex vivo* under different conditions, with or without TCR stimulation and IL-12. All conditions are with continued IL-7 and IL-15 except where indicated. Blue and light blue represent NM and NEM samples, respectively. (**B**) *Tcf7*-YFPxP14 cells were sorted on day 3 as described for (Fig. 4A) and recultured *ex vivo* in the presence of IL-2, IL-7, and IL-15 for 10 days. (**C**) *Tcf7*-YFP levels in blood and inguinal lymph nodes at days 4 and 10 after transfer. Clear histograms represent $CD45.1^+$ host cells and filled histograms represent $CD45.2^+$ donor cells. Quantification of percentage of *Tcf7*-$YFP^+$ cells in each tissue at each time point. (**D**) Representative gating strategy for *ex vivo* and *in vivo* experiments. [A-B] Mean ± s.d., data are from a single experiment. [C] Data are from a single experiment with n=2-4 biological replicates. Statistical analysis performed using two-tailed unpaired t tests between *Tcf7* low and *Tcf7* high cells for each tissue at each time point. **$p<0.01$.

Figure S5

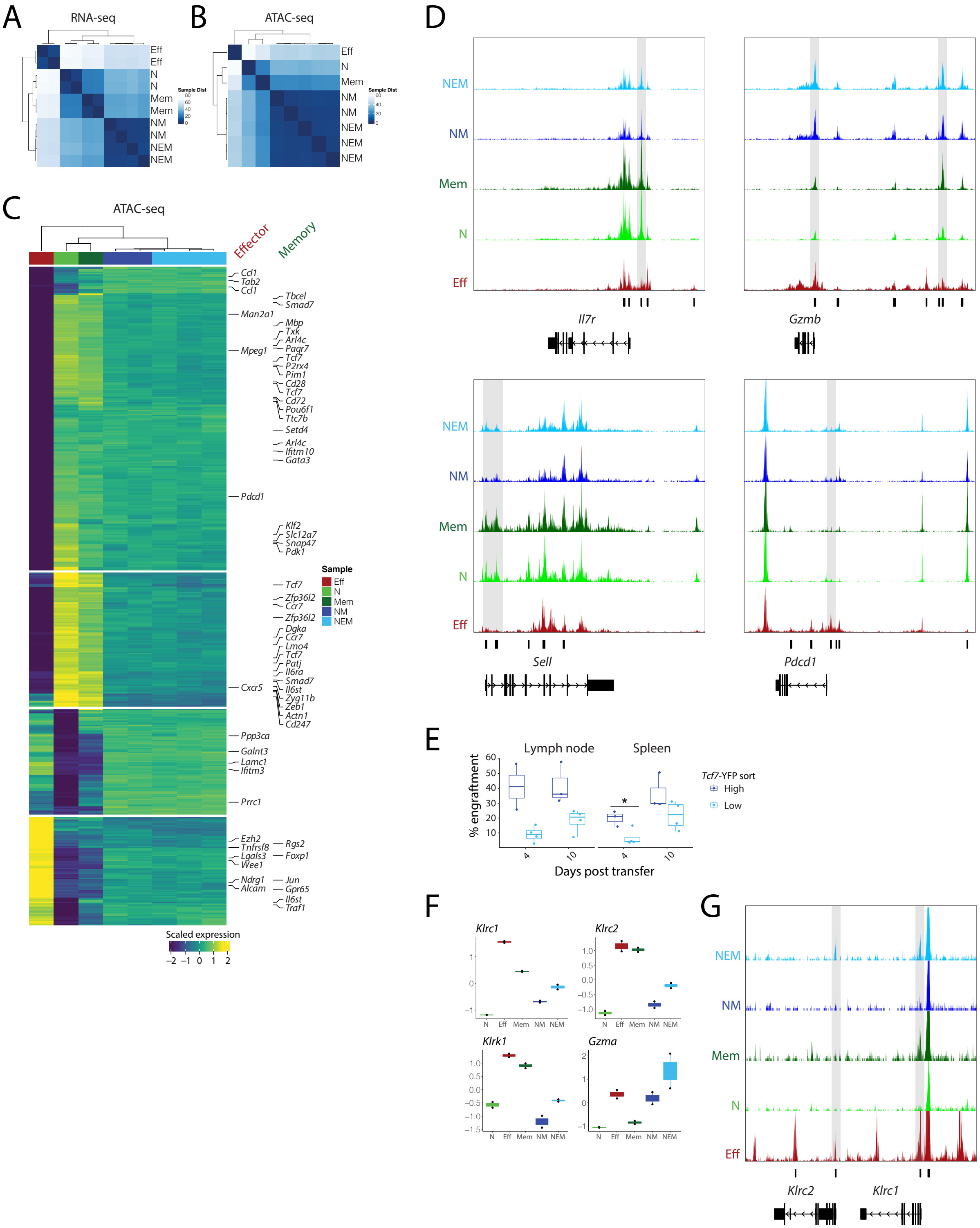

A
RNA-seq
B
ATAC-seq
C
ATAC-seq
Effector
Memory
Sample
Eff
N
Mem
NM
NEM
Scaled expression
D
Il7r
Gzmb
Sell
Pdcd1
E
Lymph node
Spleen
% engraftment
Days post transfer
Tcf7-YFP sort
High
Low
F
Klrc1
Klrc2
Klrk1
Gzma
G
Klrc2
Klrc1

**Figure S5: Memory programming in cells that decide to maintain *Tcf7* expression early (NM) or reactivate *Tcf7* later (NEM).** (**A**) Alternative representation to Fig. 5A: correlation matrix of RNA-seq profiles (top 500 DEG) for recultured cells compared to effector (Eff) and day 0 naive (N) and memory (Mem) controls. (**B**) Alternative representation to Fig. 5F: PCA of ATAC-seq counts of top 500 differential peaks between recultured cells and controls. (**C**) Heatmap displaying top 500 differentially accessible peaks (lfc $\geq 2$, Bonferroni-adjusted p value $< 0.05$) between recultured populations and Eff, N, and Mem controls. Color legend indicates row z-scores of regularized log transformed count data. Memory and effector associated genes from MSigDB Goldrath and Kaech collections are highlighted in green and red, respectively. (**D**) ATAC-seq read coverage tracks. Vertical bars annotate differentially accessible peaks between recultured cells and controls, and grey shading highlights peaks of interest. (**E**) Engraftment of *Tcf7*-YFP sorted low and high cells in secondary lymphoid organs after adoptive transfer (see Fig. 4A), quantified as the percentage of transferred CD45.2$^+$ cells in each organ divided by the percentage of CD45.2$^+$ cells in the blood. (**F**) Scaled gene expression from bulk RNA-seq for DE genes (lfc $\geq 2$, Bonferroni-adjusted p value $< 0.05$ except for *Gzma*: non-adjusted p = 0.0175 ) between D9 recultured NM and NEM cells (see Fig. 5). (**G**) ATAC-seq read coverage tracks. Vertical bars annotate differentially accessible peaks between recultured cells and controls, and grey shading highlights peaks of interest.[A, F] n = 2 biological replicates for each sample. [B-D, G] n = 1 biological replicate for Eff, N, Mem, n = 2 for NM, n = 3 for NEM. [E] Data are from a single experiment with n=2-4 biological replicates. Statistical analysis performed using two-tailed unpaired t tests between *Tcf7* low and *Tcf7* high cells for each tissue at each time point. *p<0.05.

Figure S6

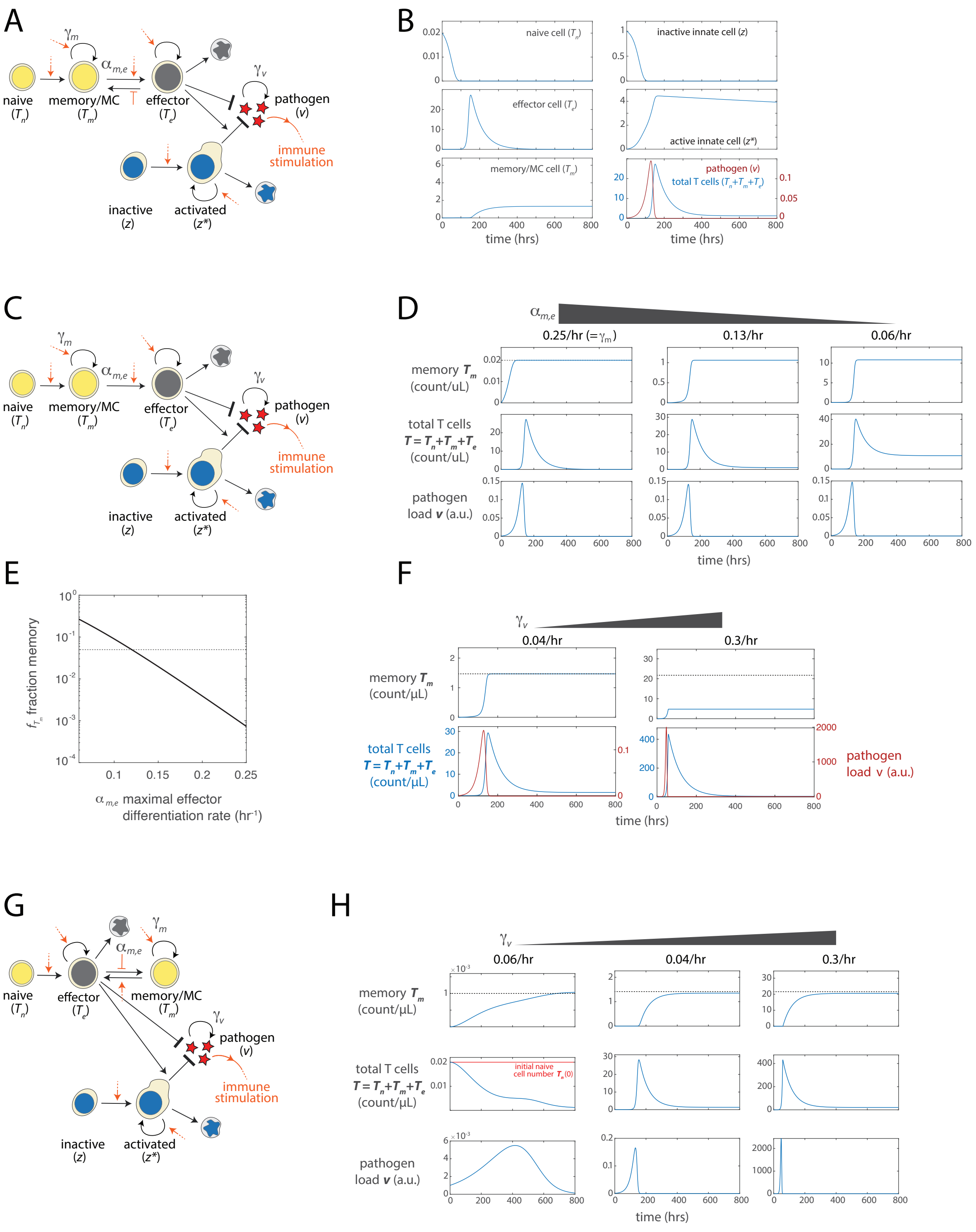

A
naive ($T_n$)
memory/MC ($T_m$)
effector ($T_e$)
$\gamma_m$
$\alpha_{m,e}$
$\gamma_v$
pathogen ($v$)
immune stimulation
inactive ($z$)
activated ($z^*$)
B
naive cell ($T_n$)
inactive innate cell ($z$)
effector cell ($T_e$)
active innate cell ($z^*$)
memory/MC cell ($T_m$)
pathogen ($v$)
total T cells ($T_n+T_m+T_e$)
time (hrs)
C
naive ($T_n$)
memory/MC ($T_m$)
effector ($T_e$)
pathogen ($v$)
immune stimulation
inactive ($z$)
activated ($z^*$)
D
$\alpha_{m,e}$
0.25/hr (=$\gamma_m$)
0.13/hr
0.06/hr
memory $T_m$ (count/uL)
total T cells $T=T_n+T_m+T_e$ (count/uL)
pathogen load $v$ (a.u.)
time (hrs)
E
$f_{T_m}$ fraction memory
$\alpha_{m,e}$ maximal effector differentiation rate (hr$^{-1}$)
F
$\gamma_v$
0.04/hr
0.3/hr
memory $T_m$ (count/µL)
total T cells $T=T_n+T_m+T_e$ (count/µL)
pathogen load v (a.u.)
time (hrs)
G
naive ($T_n$)
effector ($T_e$)
memory/MC ($T_m$)
$\alpha_{m,e}$
$\gamma_m$
$\gamma_v$
pathogen ($v$)
immune stimulation
inactive ($z$)
activated ($z^*$)
H
$\gamma_v$
0.06/hr
0.04/hr
0.3/hr
memory $T_m$ (count/µL)
total T cells $T=T_n+T_m+T_e$ (count/µL)
initial naive cell number $T_n(0)$
pathogen load $v$ (a.u.)
time (hrs)

**Figure S6: Immune response dynamics for flexible, early, and late decision models.**

*The flexible T cell decision making model (A-B) recapitulates the canonical immune response.* (**A**) Schematic for the flexible decision model, including the rate constants for effector differentiation ($\alpha_{m,e}$), memory precursor proliferation ($\gamma_m$), and pathogen replication ($\gamma_v$). (**B**) Time traces show the time evolution of the levels of naive T cells ($T_n$), effector T cells ($T_e$), memory/MC T cells ($T_m$), inactive innate immune cells ($z$), active innate immune cells ($z^*$), and pathogen load ($v$). Following pathogen emergence, there is an increase in the numbers of activated innate immune cells, followed by an expansion in T cell number that occurs concomitantly with a decrease in pathogen load. Following pathogen clearance, there is a decline in T cell numbers to a stable level, reflecting a rise in the number of long-lived memory cells. *In the early (irreversible) decision model (C-F), the balance between effector differentiation and memory precursor proliferation sets the size of the memory population.* (**C**) Schematic for the early decision model, indicating the rate constants for effector differentiation ($\alpha_{m,e}$), memory precursor proliferation ($\gamma_m$), and pathogen replication ($\gamma_v$). (**D**) Simulation traces showing time evolution of memory cell numbers (top), total T cell numbers (middle), and pathogen load (bottom, $v$), obtained for different values of $\alpha_{m,e}$. Dotted line (top left) shows the initial number of naive cells in the system. Reducing the rate of effector differentiation relative to memory precursor proliferation increases the size of the resultant memory population. (**E**) Plot showing the fraction of cells at the peak of expansion that become memory cells, $f_{Tm}$, as a function of the effector differentiation rate $\alpha_{m,e}$. Dotted line shows a memory fraction $f_{Tm} = 0.05$, used to set the differentiation rate for comparison with the reversible switching model. (**F**) Simulation traces showing time evolution of memory cell numbers (top), total T cell numbers and pathogen load (bottom), for different pathogen replication rates. *A late (obligate reversible) decision strategy*

*(G-H) enables scalable memory generation but fails to generate memory upon a mild challenge.* (**G**) Schematic for the late decision model, indicating the rate constants for effector differentiation ($\alpha_{m,e}$), memory precursor proliferation ($\gamma_m$), and pathogen replication ($\gamma_v$). (**H**) Simulation traces showing time evolution of memory cell numbers (top), total T cell numbers (middle), and pathogen load (bottom, $v$), obtained for different rates of pathogen replication $\gamma_v$. Dotted line (top) shows the final memory cell number in the system. Red line (middle left) indicates the initial number of naive cells in the system. At low rates of pathogen proliferation, the number of memory cells formed are only a small fraction of the initial number of naive cells; the system is unable to store substantial amounts of memory in this regime. See Mathematical Appendix Table 2 for parameter values used for simulations of all models.

### Mathematical Appendix

#### Contents

**1 Introduction** **13**

**2 Models of memory formation** **14**

### 1 Introduction

The immune system remembers prior pathogen encounters through the production of memory B- and T cells during an acute infection. It stores information about the nature and severity of the encountered pathogen, through the number, antigen specificity and functional states of the memory cells that form. The quantity of T cells generated, in particular, encodes information of the severity of a prior infection; across a range of pathogenic challenges, it has been observed that the number of memory cells that arise is a fixed fraction of the total number of T cells at the peak of the infection[3–5]. This linear scaling in memory production allows the body to generate a proportionally larger quantity of memory when faced with a more severe pathogenic challenge.

This scalable control of memory cell generation may reflect a consequence of the fate decisions made by responding T cells during the course of infection. In the face of stimulatory signals, T cells decide whether to differentiate into a short-lived effector cell, or to maintain self-renewal potential and become a memory cell once the pathogen is cleared. Through our single-cell tracking studies, we have identified a strategy, whereby T cells initially retain memory potential shortly after activation, but stochastically transition into an effector state with a probability that increases with the strength of antigen stimulation. Importantly, the decision to become an effector cell is flexible, and this effector dedifferentiation occurs when antigenic signals are no longer present.

Here, we analyze the population dynamics of CD8 T cells to characterize memory formation using this flexible decision making strategy. We specifically evaluate whether this strategy permits quantitative encoding of infection severity through such scalable modulation of memory population sizes. To do so, we develop a mathematical framework to model the immune response to acute infection, that incorporates pathogen replication, control by effector T cells, as well as the flexible decision making strategy discussed above ("Flexible decision model"). Additionally, our model explicitly accounts for activity of the innate immune system, which acts both as a first-line responder and also as a executor of T cell-directed effector activity. We further compare the memory encoding capabilities of the flexible strategy with two other decision making strategies proposed in the literature: one where all memory cells arise early, as a result of irreversibility in the decision to become short-lived effector cells ("Early decision model")[6–8]; and another, where all memory cells arise from effector cells, as a result of a direct conversion of naive cells to effector cells following antigen encounter ("Late decision model")[9–11]. By comparing the performance of these different models using modeling, we aim to gain insight as to why certain decision-making strategies may have been functionally beneficial for pathogen defense by the immune system and hence, selected for during evolution.

### 2 Models of memory formation

#### Flexible decision model

Here, we describe the reversible model for T cell memory decision making (Fig. 6). In this model, naive cells first transition into memory cell precursors in the presence of pathogens. These memory precursors then transition into effector cells with a probability that increases with increasing pathogen load. Both memory precursors and effector cells proliferate with a similar rate that increases with antigen level, as previously observed[12]; however, only effector cells undergo apoptosis as a result of activation-induced cell death, a reflection of their short-lived nature. In addition to T cells, we also consider the innate immune system in both its inactive and active states, and collectively model the innate immune response in these two states using two variables. T cells and innate cells then mediate pathogen killing, both independently from each other and also in a cooperative manner. This model is described by the following system of equations:

$$\begin{aligned}
\text{naive T cell:} \quad & \frac{\mathrm{d}T_n}{\mathrm{d}t} = -\alpha_n v T_n \\
\text{memory T cell:} \quad & \frac{\mathrm{d}T_m}{\mathrm{d}t} = \alpha_n v T_n + \big(\frac{v}{v+K_m}\big)\,\gamma_m T_m - \big(\frac{v}{v+K_{m,e}}\big)\,\alpha_{m,e} T_m + \big(\frac{K_{e,m}}{v+K_{e,m}}\big)\,\beta_{e,m} T_e \\
\text{effector T cell:} \quad & \frac{\mathrm{d}T_e}{\mathrm{d}t} = \big(\frac{v}{v+K_e}\big)\,\gamma_e T_e + \big(\frac{v}{v+K_{m,e}}\big)\,\alpha_{m,e} T_m - \big(\frac{K_{e,m}}{v+K_{e,m}}\big)\,\beta_{e,m} T_e - \delta_e T_e \\
\text{pathogen:} \quad & \frac{\mathrm{d}v}{\mathrm{d}t} = \big(\frac{v}{\epsilon+v}\big)^N \gamma_v v - \big(\delta_{v_1} T_e + \delta_{v_2} z^* + \delta_{v_3} z^* \, T_e\big)\, v \\
\text{inactive innate cell:} \quad & \frac{\mathrm{d}z}{\mathrm{d}t} = -\alpha_z v \, z \\
\text{active innate cell:} \quad & \frac{\mathrm{d}z^*}{\mathrm{d}t} = \alpha_z v \, z + \big(\frac{v}{v+K_{z^*}}\big)\,\gamma_{z^*} \, z^* - \delta_{z^*} \, z^* \qquad (1)
\end{aligned}$$

Here, the subscripts $n$, $m$, and $e$ denote naive, memory, and effector types, respectively, $v$ denotes the pathogen population, and $z$ and $z^*$ denote the inactive and active innate immune cells, respectively. The rates $\alpha_{x,y}$ denote differentiation rates from the $x$ to the $y$ cell type (for $x, y \in \{n, m, e\}$), $\beta_{e,m}$ denotes the de-differentiation rate from effector to memory (i.e., the only reversed differentiation in the model), $\gamma_x$ denotes proliferation rate of cell type $x$, and $\delta_x$ denotes the death rate of cell type $x$. The parameters $K_{(\cdot)}$ denote the pathogen load for half maximal rate of a process indicated in the subscript.

A detailed description of the model variables, parameters and initial conditions are given in Tables 1, 2. Parameters have been chosen based on the immune compartment sizes as measured in mice, as well as T cell biological parameters that we and others have measured. The initial conditions have been chosen to reflect the initial onset of an infection by a pathogen for which no prior immunological memory has been developed (Table 1); specifically, antigen-specific naive cells are present at low amounts, effector and memory cells are absent and the initiating pathogen is introduced at a low initial level.

| | Variable description | Initial value |
|---|---|---|
| $T_n$ | naive T cell | 0.02 cells/µL |
| $T_m$ | memory T cell | 0 cells/µL |
| $T_e$ | effector T cell | 0 cells/µL |
| $v$ | pathogen load | $10^{-3}$ units/µL |
| $z$ | inactive innate cell | 1 cell/µL |
| $z^*$ | activated innate cell | 0 cells/µL |

Table 1: **Variables and initial conditions for the reversible epigenetic switch model**

| | Parameter description | Value |
|---|---|---|
| $\alpha_n$ | naive cell activation rate constant | 5/(units/µL)/hr |
| $\gamma_m$ | maximal memory cell proliferation rate constant | 0.25/hr |
| $K_m$ | pathogen load for half maximal memory cell proliferation | 0.1 units/µL |
| $\alpha_{m,e}$ | maximal effector differentiation rate constant | 0.25/hr |
| $K_{m,e}$ | pathogen load for half maximal effector differentiation | 0.1 units/µL |
| $\gamma_e$ | maximal effector cell proliferation rate constant | 0.25/hr |
| $K_e$ | pathogen load for half maximal effector proliferation | 0.1 units/µL |
| $\beta_{e,m}$ | maximal rate of effector de-differentiation | $8\times10^{-4}$/hr |
| $K_{e,m}$ | pathogen load for half maximal effector de-differentiation | $2.5\times10^{-3}$ units/µL |
| $\delta_e$ | rate constant for effector cell death | 0.016/hr |
| $\gamma_v$ | rate of pathogen replication | 0.003-0.45/hr |
| $\epsilon$ | pathogen load for extinction | $10^{-4}$ /µL |
| $N$ | sharpness of extinction effect for pathogen | 100 |
| $\delta_{v_1}$ | rate constant for T cell pathogen killing | $4.5\times10^{-3}$/(cells/µL)/hr |
| $\delta_{v_2}$ | rate constant for innate cell pathogen killing | $1.5\times10^{-3}$/(cells/µL)/hr |
| $\delta_{v_3}$ | rate constant for T-cell assisted innate cell pathogen killing | $1.5\times10^{-3}$/(cells/µL)$^{-2}$/hr |
| $\alpha_z$ | innate cell activation rate constant | 2.5/(units/µL)/hr |
| $\gamma_{z^*}$ | maximal activated innate cell proliferation rate | 0.02/hr |
| $K_{z^*}$ | pathogen load for half maximal innate cell proliferation | 0.01 units/µL |
| $\delta_{z^*}$ | turnover rate for activated innate cell | $2\times10^{-4}/hr$ |

Table 2: **Parameters for the reversible epigenetic switch model**

We point out that this system is fundamentally a Lotka-Volterra model where immune cells are predators and pathogens are prey. However, we have modified this framework to describe the immune response in the following ways: first, we have incorporated saturation terms in the rates of pathogen-induced T cell and innate cell division ($K_m$, $K_e$), as well as T cell effector to memory differentiation ($K_{m,e}$) and de-differentiation ($K_{e,m}$). The values for these saturation terms are further chosen to reflect the biological upper-bounds for these cellular processes. Second, we incorporate a threshold pathogen load, $\epsilon$, below which the pathogen replication rate drops to zero, reflecting extinction of the pathogen. This pathogen extinction threshold ensures that this deterministic system of equations, when simulated, has a well-defined behavior and is not subject to numerical integration errors at very low pathogen loads[13]. However, we note that the dynamics of simulated response does not generally depend on the exact value of the extinction coefficient chosen.

From numerical simulations, we see that the flexible decision model reproduces the canonical dynamics of the adaptive immune response (Fig. 6B, Fig. S6A-B). Upon introduction to the system, pathogens increase exponentially in number, giving rise to a subsequent expansion of the T cell numbers from their initial low levels in the naive cell population. This expansion occurs concomitantly with pathogen clearance, and is followed by a decline in T cell numbers to a stable elevated baseline, reflecting the generation of long-lived memory cells that can survive following pathogen clearance. During the course of the immune response, the numbers of activated innate immune cells increases rapidly and decreases steadily for the reminder of simulation (Fig. S6A-B). This heightened innate immune activity is critical for ensuring that pathogens clear after T cell contraction and do not rebound in number.

How does the size of the generated memory population depend on the severity of infection in the reversible switching model? In particular, we wish to ascertain whether this system can produce memory cells in numbers that scale linearly with the peak T cell numbers during an infection, as observed experimentally[1]. To ask this question, we performed simulations of the system with different values of pathogen replication rate, $\gamma_v$, as a means to vary pathogen virulence. We found that pathogens with different replication rates gave rise to different degrees of T cell expansion and contraction, with faster-replicating pathogens generating a stronger T cell response, as expected (Fig. 6B).

However, the fraction $f_{T_m}$ of memory cells to the total number of T cells present at the expansion peak becomes a fixed number (Fig. 6B, 6C: shaded area). Specifically, in the regime where there is substantial T cell expansion ($> 10^2$ fold relative to naive cell numbers), the memory fraction $f_{T_m}$ remains constant for a broad range of viral replication rates $\gamma_v$, spanning an order of magnitude (Fig. 6C, top, shaded area). On the other hand, for slowly growing pathogens (small $\gamma_v$), the memory fraction $f_{T_m}$ increases, with a non-linear inverse dependence on the virulence and the average viral load accumulated during the infection (Fig. 6C); in this regime, the number of memory cells depends strongly on the initial number of naive cells present. In summary, these results show that a flexible switching strategy for T cell memory generation allows for the amount of the generated T cell memory to scale with the size of the T cell response, in a way that depends on the severity of the infection. Our analytical results in the following section well recapitulates the behavior of this memory fraction, as indicated in Fig. 6.

##### Analytical results for the flexible decision model

Consider the dynamics of memory $T_m$ and effector $T_e$ populations, given by eq. 1. We begin by identifying the dominant processes in different regimes of accumulated viral load $v$.

Specifically, we compare the viral load with the half maximal loads ($K_{(\cdot)}$'s in eq. 1), necessary for different processes. Below the terms that are relevant for the dynamics of memory and effetor populations in the high viral load ($v \gg K_m, K_{m,e}$) and the low viral load ($v \ll K_{e,m}$) regimes are indicated:

$$\text{memory T cell:} \qquad \frac{\mathrm{d}T_m}{\mathrm{d}t} = \underbrace{\alpha_n v T_n + (\frac{v}{v+K_m}) \cdot \gamma_m T_m - (\frac{v}{v+K_{m,e}}) \cdot \alpha_{m,e} T_m}_{v \gg K_m, K_{m,e}} + \underbrace{(\frac{K_{e,m}}{v+K_{e,m}}) \cdot \beta_{e,m} T_e}_{v \ll K_{e,m}} \tag{2}$$

$$\text{effector T cell:} \qquad \frac{\mathrm{d}T_e}{\mathrm{d}t} = \underbrace{(\frac{v}{v+K_e}) \cdot \gamma_e T_e + (\frac{v}{v+K_{m,e}}) \cdot \alpha_{m,e} T_m}_{v \gg K_e, K_{m,e}} - \underbrace{(\frac{K_{e,m}}{v+K_{e,m}}) \cdot \beta_{e,m} T_e}_{v \ll K_{e,m}} - \delta_e T_e. \tag{3}$$

We can formally integrate over the dynamical equation in eq. 2 to find a formal solution for the number of memory cells $T_m(t)$ at time $t$ post infection,

$$\begin{aligned} T_m(t) &= T_m(0) \exp\left[\int_0^t \gamma_m - \alpha_{m,e} - \frac{\gamma_m K_m}{v(s)+K_m} + \frac{\alpha_m K_{m,e}}{v(s)+K_{m,e}} \mathrm{d}s\right] \\ &\quad + \left[\int_0^t \exp\left[\int_s^t \gamma_m - \alpha_{m,e} - \frac{\gamma_m K_m}{v(s)+K_m} + \frac{\alpha_m K_{m,e}}{v(s)+K_{m,e}} \mathrm{d}s\right] \right. \\ &\quad \left. \times \left(\alpha_n v(s) T_n(s) + \frac{K_{e,m}}{v(s)+K_{e,m}} \beta_{e,m} T_e(s)\right) \mathrm{d}s \right] \end{aligned} \tag{4}$$

$$\begin{aligned} &\approx T_m(0) \exp\left[(\gamma_m - \alpha_{m,e}) \min(t, \tau_{v\approx 0})\right] \\ &\quad + \int_0^{\min(t,\tau_{v\approx 0})} \exp\left[(\gamma_m - \alpha_{m,e})(\min(t, \tau_{v\approx 0}) - s)\right] \alpha_n v(s) T_n(s) \mathrm{d}s \\ &\quad + H(t - \tau_{v\approx 0}) \int_{\tau_{v\approx 0}}^t \beta_{e,m} T_e(s) \mathrm{d}s, \end{aligned} \tag{5}$$

where $\tau_{v\approx 0}$ is the time to effectively clear the infection, and $H(t - \tau_{v\approx 0})$ is a heaviside step function that takes value 1 for $t > \tau_{v\approx 0}$, and 0, otherwise. In arriving at eq. 5, we assumed that the typical viral load over the course of the infection is much higher than the differentiation thresholds $K_m, K_{m,e}, K_e$, and $K_{e,m}$, and thus, we approximated these processes by their maximal rates in eq. 4.

The following terms are important in determining the size of the memory pool:

1. $T_m(0)$: the initial memory size;
2. $\gamma_m - \alpha_{m,e}$: the effective growth rate of the memory pool;
3. $\tau_{v\approx 0}$: time to effectively clear the infection;

4. $\beta_{e,m}$: the transition rate from effector to memory.

Here, we are interested in an immune response to a primary infection, and therefore, we can assume that $T_m(0) = 0$. Moreover, given the rates reported in Table 2, we can neglect the effective growth of the memory pool, i.e., $\gamma_m - \alpha_{m,e} \approx 0$. With these assumptions, the size of the memory pool from eq. 5 follows,

$$T_m(t) \approx \int_0^{\min(t,\tau_{v\approx 0})} \alpha_n v(s) T_n(s) ds + H(t - \tau_{v\approx 0}) \int_{\tau_{v\approx 0}}^{t} \beta_{e,m} T_e(s) ds.$$

Our goal is to estimate the asymptotic (long-term) fraction of memory to the total number of T cells (primarily effector cells) present at the expansion peak $f_{T_m} = T_m(\infty)/T_e^{\max}$. The asymptotic amount of memory follows,

$$T_m(t \to \infty) \approx \int_0^{\tau_{v\approx 0}} \alpha_n v(s) T_n(s) \mathrm{d}s + \int_{\tau_{v\approx 0}}^{\infty} \beta_{e,m} T_e(s) \mathrm{d}s,$$

then assuming $\tau_{v\approx 0} \approx \tau_{e^{\max}}$,

$$\begin{aligned} T_m(\infty) &\approx \int_0^{\tau_{e^{\max}}} \alpha_n v(s) T_n(s) \mathrm{d}s + \int_{\tau_{e^{\max}}}^{\infty} \beta_{e,m} T_e(s) \mathrm{d}s \\ &\approx \int_0^{\tau_{e^{\max}}} \alpha_n v(s) T_n(s) \mathrm{d}s + \int_{\tau_{e^{\max}}}^{\infty} \beta_{e,m} T_e^{\max} e^{-(\beta_{e,m}+\delta_e)s} \mathrm{d}s \\ &\approx \int_0^{\tau_{e^{\max}}} \alpha_n v(s) T_n(s) \mathrm{d}s + \beta_{e,m} \frac{T_e^{\max}}{\beta_{e,m} + \delta_e} \end{aligned} \tag{6}$$

where we used the relation $T_e(t > \tau_{e^{\max}}) = T_e e^{-(\beta_{e,m}+\delta_e)s}$, indicating an exponential decay of effector cells after the peak of the response $(v \ll K_{e,m})$ , from eq. 3.

From eq. 1, we can also formally express the size of the naive pool as,

$$T_n(t) = T_n(0) \exp\left[-\alpha_n \int_0^t v(s) ds\right],$$

Therefore, the first term in the solution of eq. 6 follows,

$$\alpha_n \int_0^{\tau_{e^{\max}}} v(s) T_n(s) ds = T_n(0) \left[1 - e^{-\alpha_n \tilde{V}}\right], \quad \text{with} \quad \tilde{V} = \int_0^{\tau_{e^{\max}}} v(r) \mathrm{d}r$$

Here, $\tilde{V}$ reflects the total amount of pathogens accumulated during the infection. Putting it all together, we find

$$T_m(\infty) \approx T_n(0) \left[1 - e^{-\alpha_n \tilde{V}}\right] + \beta_{e,m} \frac{T_e^{\max}}{\beta_{e,m} + \delta_e}, \tag{7}$$

resulting the following memory fraction,

$$f_{T_m} = \frac{T_m(\infty)}{T_e^{\max}} \approx \frac{x(0)}{T_e^{\max}} \left[1 - e^{-\alpha_n \tilde{V}}\right] + \frac{\beta_{e,m}}{\beta_{e,m} + \delta_e}. \tag{8}$$

When $T_e^{\max} \gg T_n(0)$, we recover the constant memory fraction $f_{T_m} \approx \frac{\beta_{e,m}}{\beta_{e,m}+\delta_e}$.

So far we have assumed that $\gamma_m - \alpha_{m,e} \approx 0$. When this assumption does not hold, the first term in our expression for $f_{T_m}$ becomes,

$$
\begin{aligned}
C_0 &= \frac{1}{T_e^{\max}} \int_0^{\tau_{e^{\max}}} \exp\left[(\gamma_m - \alpha_{m,e})(\tau_{e^{\max}} - s)\right] \alpha_n v(s) T_n(s) \mathrm{d}s \\
&= \frac{\alpha_n T_n(0)}{T_e^{\max}} \int_0^{\tau_{e^{\max}}} \exp\left[(\gamma_m - \alpha_{m,e})(\tau_{e^{\max}} - s) - \alpha_n \int_0^s v(r)\mathrm{d}r\right] v(s)\mathrm{d}s \\
&\leq \frac{\alpha_n T_n(0)}{T_e^{\max}} \int_0^{\tau_{e^{\max}}} \exp\left[(\gamma_m - \alpha_{m,e})(\tau_{e^{\max}} - s)\right] v(s)\mathrm{d}s,
\end{aligned}
\tag{9}
$$

with a strong dependence on $\gamma_m - \alpha_{m,e}$.

Note that the innate immune dynamics does not explicitly determine the memory fraction $f_{T_m}$, however, it influences the magnitude of $\tilde{V}$ and so it is expected to be important in the low viral replication $\gamma_v$ regime.

#### Early decision model

From experimental studies, it has been proposed that memory cells originate primarily from cells that have undergone little or no effector differentiation, and that memory precursors, upon silencing the memory regulator TCF1 and differentiating, are committed to becoming short-lived effectors[6,14]. Using the mathematical modeling framework developed above, we evaluate whether this irreversible effector decision strategy could also enable the asymptotic (long-term) memory T cell numbers $T_m(\infty)$ to scale linearly with the peak T cell number (primary effector cells) $T_e^{\max}$. To do so, we performed simulations of the above model, rendering effector differentiation irreversible by setting the rate of effector de-differentiation $\beta_{e,m}$ to zero.

From our simulations, we found that the number of memory T cells emerging depends on the balance between effector differentiation and memory precursor proliferation, as determined by the rate constants $\alpha_{m,e}$ and $\gamma_m$ respectively. When these two rate constants are equal, the number of generated memory cells cannot exceed the initial number of naive cells (Fig. S6C-F). This is because the memory cell population, upon emerging from the naive cell pool, cannot further change in number as proliferation is balanced exactly by differentiation. We note that this regime captures the dynamics of obligate asymmetric division, where the division of each memory precursor necessarily gives rise to a precursor and a differentiated progeny. When the rate of effector differentiation $\alpha_{m,e}$ is smaller than that of memory precursor proliferation $\gamma_m$, the number of memory cells can exceed the initial naive cell number, due to a net proliferation of this population; in this regime, the size of the memory pool grows with increasing $\gamma_v$.

To perform a mathematically comparable comparison of the flexible and irreversible switching models[15], we chose a rate of effector differentiation for the latter model to be $\alpha_{m,e} = 0.12/\text{hr}$, such that the fraction of memory cells generated under conditions of moderate pathogen virulence ($\gamma_m = 0.04/\text{hr}$) were equivalent for the two models with $f_{T_m} = 0.05$.

All other parameters were kept constant. These simulations show that the irreversible switching model is unable to generate a constant fraction of memory cells amid changes in pathogen replication rates (Fig. 6C middle). The memory fraction was upheld at a similar value $f_{T_m} = 0.05$ at moderate pathogen replication rates $\gamma_v = 0.04/hr$; however, this fraction decreased steadily with increasing $\gamma_v$, eventually approaching less than 0.01 at high pathogen replication rates (Fig. 6C, middle; Fig. S6C-F). Indeed, from an approximate analytical solution of this system, we found that the memory cell fraction has an inherently inverse dependence on the peak T cell population size, as follows:

$$f_{T_m} \approx \frac{T_n(0)}{T_e^{\max}} \left[1 - e^{-\alpha_n \tilde{V}}\right] \tag{10}$$

This dependence cannot be offset when the fraction is much smaller than unity, which typically holds in the regime where the majority of cells generated at the height of an acute infection are those with effector function. Thus, from these results, we conclude that the T cell decision making strategy, where the memory precursors switch irreversibly into becoming short-lived effectors, cannot produce memory T cell numbers that scale proportionally with the peak T cell population sizes.

#### Late decision model (obligate reversible decision model)

An alternate strategy for memory differentiation is that, upon activation, all naive cells must first pass through an effector stage prior to the decision as whether to differentiate into memory (Fig. 6, bottom). This view is supported by the evidence that cells with a history of effector gene expression can become memory cells[10], and that cells on the road to forming memory retain chromatin signatures of the effector state, while harboring the ability to re-activate memory genes that are silenced during effector differentiation[9]; this model can be thought as an obligate reversible decision model.

To evaluate such a decision making strategy, we alter our model above, such that naive cells, upon activation, directly transition to an effector state instead of a memory precursor state (Fig. 6, bottom, Fig. S6G-H). Following the model in eq. 1, the ordinary differential equations describing the T cell populations in the late decision model are modified as follows:

$$\begin{aligned}
\text{naive T cell:} \quad \frac{\mathrm{d}T_n}{\mathrm{d}t} &= -\alpha_n v T_n \\
\text{memory T cell:} \quad \frac{\mathrm{d}T_m}{\mathrm{d}t} &= \Big(\frac{v}{v+K_m}\Big) \cdot \gamma_m T_m - \Big(\frac{v}{v+K_{m,e}}\Big) \cdot \alpha_{m,e} T_m + \Big(\frac{K_e}{v+K_e}\Big) \cdot \beta_{e,m} T_e \\
\text{effector T cell:} \quad \frac{\mathrm{d}T_e}{\mathrm{d}t} &= \alpha_n v T_n + \Big(\frac{v}{v+K_e}\Big) \cdot \gamma_e T_e + \Big(\frac{v}{v+K_{m,e}}\Big) \cdot \alpha_{m,e} T_m \\
&\quad - \Big(\frac{K_{e,m}}{v+K_{e,m}}\Big) \cdot \beta_{e,m} T_e - \delta_e T_e
\end{aligned}$$

For direct comparison of this decision making strategy to the model with flexible switching, we keep all the parameters unchanged. From simulations, we find this obligate reversible switching strategy can generate constant fractions of memory cells over a range of pathogen proliferation rates, but fails to generate any substantial memory when pathogens replicate slowly (small $\gamma_v$) and the ensuing immune responses are mild. When pathogens replicate rapidly and give rise to a substantial T cell expansion, memory cells form robustly at defined fraction and number, similar to the flexible switching model (Fig. 6C, bottom, Fig. S6G-H, center, right); however, when pathogens proliferate very slowly, such that there is minimal amount of T cell expansion, the number of formed memory cells constitutes only a small fraction of the starting naive cells. Consequently, in this regime, the ability of the immune system to respond to a secondary challenge is no longer heightened, and is likely compromised.

##### Analysis of late decision model

Following the analytical analyses for the flexible switching model, we can again identify the dominant terms in different regimes of viral load. By assuming that the viral load triggered by the infection is much larger than the cellular differentiation thresholds $K_m, K_{m,e}, K_e$, and $K_{e,m}$, we arrive at the following approximate expression for the size of the memory pool $T_m(t)$ at time $t$ in the late decision model.

$$T_m(t) \approx T_m(0) \exp\left[(\gamma_m - \alpha_{m,e}) \min(t, \tau_{v\approx 0})\right] + H(t - \tau_{v\approx 0}) \int_{\tau_{v\approx 0}}^{t} \beta_{e,m} T_e(s) \mathrm{d}s.$$

Then for primary immune response ($T_m(0) = 0$) we have,

$$f_{T_m} = \frac{T_m(\infty)}{T_e^{\max}} \approx \frac{\beta_{e,m}}{\beta_{e,m} + \delta_e}. \tag{11}$$

We note that this expression does not hold in the regime of slow/inefficient virus dynamics, consistent with results from simulations (Fig. 6).
